# MuSHIN: A multi-way SMILES-based hypergraph inference network for metabolic model reconstruction

**DOI:** 10.1101/2025.07.07.663526

**Authors:** Yanlong Zhao, Yixiao Chen, Yi Yu, Xiang Liu, Jiawen Du, Jun Wen, Chen Liao, Quan Sun, Ren Wang, Can Chen

**Author notes:** These authors contributed equally to this work (Yanlong Zhao, Yixiao Chen).

## Abstract

Genome-scale metabolic models (GEMs) are indispensable tools for probing cellular metabolism, enabling predictions of metabolic fluxes, guiding strain optimization, and advancing biomedical research. However, their predictive capacity is often compromised by incomplete reaction networks, stemming from gaps in biochemical knowledge, annotation inaccuracies, and insufficient experimental validations. Here we present MuSHIN (**Mu**lti-way **S**MILES-based **H**ypergraph **I**nterface **N**etwork), a novel deep hypergraph learning method that integrates network topology with biochemical domain knowledge to predict missing reactions in GEMs. Evaluated on 926 high- and intermediate-quality GEMs with artificially removed reactions, MuSHIN significantly outperforms state-of-the-art methods, achieving up to a 17% improvement across multiple metrics and maintaining robust recovery even under severe network sparsity. Furthermore, MuSHIN substantially enhances phenotypic predictions in 24 draft GEMs associated with fermentation by resolving critical metabolic gaps, as validated against experimental measurements. Together, these findings highlight MuSHIN’s potential to advance GEM reconstruction and accelerate discoveries in systems biology, metabolic engineering, and precision medicine.

## Introduction

Genome-scale metabolic models (GEMs) are foundational tools in systems biology, offering a comprehensive representation of an organism’s metabolic pathways at the genome scale^1–5^. By combining biochemical and physiological insights, GEMs allow researchers to simulate cellular metabolism in silico and estimate metabolic fluxes under varying conditions. These simulations inform strategies for genetic engineering and have driven advances in metabolic engineering^6,7^, drug discovery^8,9^, and synthetic biology^10,11^. Despite their broad utility, GEMs are often incomplete due to gaps in biochemical knowledge, annotation errors, and limited experimental validation^12,13^. This incompleteness undermines the functional integrity of GEMs, leading to inaccurate modeling of essential cellular phenotypes, such as biomass production, metabolite secretion, and gene essentiality^14–16^. The impact is particularly pronounced in non-model organisms, whose metabolic pathways remain poorly characterized^12,13^. Even in well-studied species, certain reactions are only active under specific environmental or cellular contexts^17,18^. Addressing these deficiencies is thus critical for improving GEM fidelity and expanding their applicability across organisms and experimental settings.

A variety of optimization-based strategies have been developed to address gaps in metabolic models, many of which rely on incorporating phenotypic information, such as growth profiles or flux distributions, to guide the insertion of missingg reactions^14,19–23^. These methods seek to reconcile model predictions with experimental observations by identifying inconsistencies and supplementing pathways to restore metabolic function^14^. However, their reliance on condition-specific experimental data restricts their use to well-characterized organisms, making them unsuitable for many microbial species that are uncultivable or poorly annotated^24^. In contrast, topology-driven methods such as GapFind/GapFill^22^ offer a phenotype-independent alternative by leveraging stoichiometric constraints to identify non-functional metabolites and proposing reactions that enable feasible flux through the network (see Supplementary Note 1). Despite circumventing the need for experimental measurements, these methods rely on linear optimization and simplified network assumptions, overlooking the complex multi-metabolite interactions fundamental to biochemical systems. As genome-scale models continue to expand in scope and complexity, such simplifications increasingly constrain their ability to reflect biological reality.

Hyperedge prediction offers a promising solution to these challenges by leveraging hypergraphs, a generalization of graphs in which hyperedges can simultaneously connect multiple nodes^25–30^. Metabolic networks can be naturally represented as hypergraphs, where each hyperedge corresponds to a biochemical reaction involving multiple metabolites acting as substrates and products^26,30,31^. Unlike conventional graph-based methods that model pairwise interactions, hypergraph-based representations preserve the higher-order connectivity inherent in metabolic reactions, enabling a more accurate and comprehensive characterization of metabolic network structures^12,31–33^. However, existing hypergraph-based machine learning methods have notable drawbacks (see Supplementary Note 1). For example, CMM^34^ and C3MM^35^ employ an integrated training and prediction process that includes all candidate reactions from a predefined reaction pool during training, limiting its scalability and requiring retraining whenever a new reaction pool is introduced. The neural network-based method NHP^36^ approximates hypergraphs using graphs to generate node features, leading to the loss of higher-order interactions. Recent methods such as CHESHIRE^12^ and CLOSEgaps^13^ lack the expressive capacity of attention-based hypergraph models, struggling to capture the complex multi-metabolite interactions inherent in metabolic reactions. More importantly, these methods do not fully exploit accessible domain knowledge, such as metabolite chemical structures, which are essential for evaluating reaction plausibility. These shortcomings underscore the pressing need for biologically grounded, domain-informed methods that preserve the hypergraph topology to more accurately recover missing reactions.

In this work, we introduce MuSHIN (**Mu**lti-way **S**MILES-based **H**ypergraph **I**nterface **N**etwork), a novel hypergraph-based deep learning method for predicting missing reactions in GEMs by integrating both topological structure and biochemical semantics. MuSHIN incorporates molecular-level information by embedding metabolite structures using transformer-based models, enriching the hypergraph representation with chemical semantics. A dynamic attention mechanism further refines metabolite and reaction features through iterative message passing, allowing MuSHIN to effectively learn the complex higher-order interactions between metabolites and reactions. Through comprehensive evaluations across multiple GEM databases, MuSHIN significantly outperforms state-of-the-art methods in both internal validation based on the recovery of synthetically removed reactions and external validation measured by the prediction of experimentally observed phenotypes. By bridging biochemical knowledge with network topology, MuSHIN offers a robust and scalable framework for automated GEM curation and advances the predictive accuracy of metabolic modeling in systems biology, metabolic engineering, and precision medicine.

## Results

### A brief overview of MuSHIN

MuSHIN is an advanced deep learning-based method designed to predict missing reactions in GEMs by integrating both topological and biochemical semantics of metabolic networks. It introduces two key innovations: (1) the use of transformer-based models, including RXNFP^37^ and ChemBERTa^38^, to initialize chemically informed features of metabolites and reactions; and (2) a dynamic attention mechanism^39^ that iteratively refines metabolite and reaction features through bidirectional node-hyperedge message passing. By jointly capturing structural connectivity and chemical context, MuSHIN offers a powerful and generalizable approach for reconstructing high-fidelity GEMs.

The metabolic network is represented as a hypergraph, where nodes denote metabolites and hyperedges correspond to biochemical reactions (**Fig. 1a-b**; see “Hypergraphs” in Methods). This formulation inherently captures the complex, higher-order interactions characteristic of metabolic reactions, compared to conventional pairwise graphs. For model training, we curate a balanced dataset comprising both positive and negative reactions, where positive samples are drawn from well-established GEMs, while negative samples are synthetically generated by perturbing known reactions to create chemically invalid alternatives, e.g., replacing one metabolite with a non-reactive compound (**Fig. 1c**; see “Negative reaction generation” in Methods). This deliberate generation strategy compels the model to discern biologically plausible reactions from implausible ones by leveraging subtle structural and topological differences. All reactions, including both positive and negative, are encoded into a hypergraph incidence matrix (**Fig. 1d**),which captures metabolite-reaction connectivity as a binary relationship, forming the basis for subsequent hypergraph learning. For simplicity, network directionality is not considered.

**Fig. 1:**
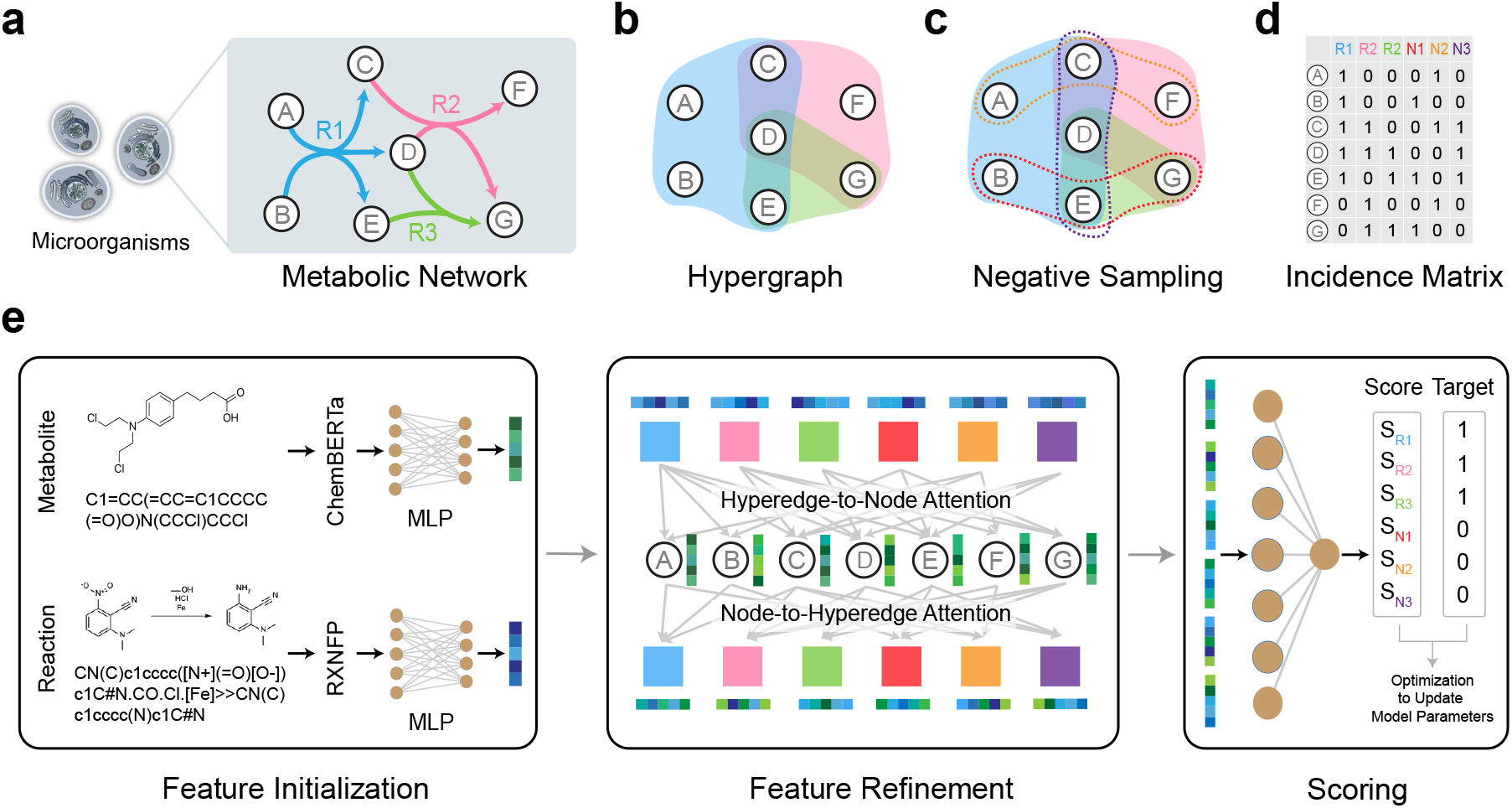
MuSHIN workflow. **a**. Schematic representation of a metabolic network. Metabolites (A-G) are nodes, and reactions (R1, R2, R3) are directed edges connecting reactants and products. **b**. Hypergraph representation of the metabolic network. Each hyperlink connects multiple metabolites participating in the same reaction. **c**. Negative sampling generates chemically invalid reactions (N1, N2, N3) by perturbing positive reactions (R1, R2, R3). Solid and dashed outlines represent positive and negative reactions, respectively. **d**. Incidence matrix of the hypergraph, encoding the binary relationships between metabolites (A-G) and reactions (R1, R2, N1, etc.). **e**. Model pipeline including: (1) feature initialization: ChemBERTa and RXNFP embed metabolites and reactions from SMILES representations, processed via MLPs to extract chemical features; (2) feature refinement: a hypergraph neural network refines node and hyperedge features through alternating hyperedge-to-node and node-to-hyperedge attention mechanisms; (3) scoring: refined reaction features are used to predict confidence scores, compared to target labels for model optimization.

The architecture of MuSHIN consists of a feature initialization stage followed by iterative feature refinement (**Fig. 1e**). During initialization, chemical information from metabolites and reactions is encoded via transformer-based models applied to their SMILES representations, a widely used textual notation that linearly encodes molecular structures. Specifically, metabolite structures are embedded using ChemBERTa^38^, while reaction SMILES are processed through RXNFP^37^, yielding rich high-dimensional feature vectors that capture detailed molecular semantics (see “Metabolite and reaction embeddings” in Methods). These vectors are then projected through multilayer perceptrons to a common dimensionality for subsequent integration. Following initialization, MuSHIN refines these features using a hypergraph neural network enhanced with a dual dynamic attention mechanism^39^. This module iteratively updates metabolite (node) and reaction (hyperedge) features via alternating node-to-hyperedge and hyperedge-to-node attention-based message passing, allowing the model to integrate higher-order dependencies and capture complex relationships within the metabolic network (see “Hypergraph neural network architecture” in Methods). Unlike conventional methods that primarily depend on pooling techniques, MuSHIN directly outputs refined context-aware reaction feature vectors, preserving nuanced local and global structural information. In the final step, the refined reaction feature vectors are fed into a single-layer neural network to predict the likelihood of reaction validity. The model is trained end-to-end by minimizing the discrepancy between predicted scores and ground-truth labels, enabling MuSHIN to faithfully capture the multifaceted structural and biochemical relationships inherent to GEMs (see Supplementary Note 2).

### MuSHIN outperforms existing methods with artificially introduced gaps

To assess the performance of MuSHIN in recovering missing reactions, we performed internal validation using GEMs with artificially introduced gaps (**Fig. 2a**, top testing scheme). Specifically, we evaluated MuSHIN against four representative baselines, including CLOSEgaps^13^, CHESHIRE^12^, NHP^36^, and HGNN^26^ (see Supplementary Note 1), across 108 models from the BiGG database^10^ and 818 models from the AGORA database^40^. Negative reactions were synthetically generated by perturbing known reactions, ensuring chemical invalidity while preserving atom balance, and paired with known valid reactions to create a balanced 1:1 positive-to-negative ratio. Each model was split into 60% training, 20% validation, and 20% testing sets. Details of the hyperparameter settings are provided in Supplementary Note 3. To ensure statistical robustness, we performed 10 Monte Carlo runs per model and reported results as the average across runs.

**Fig. 2:**
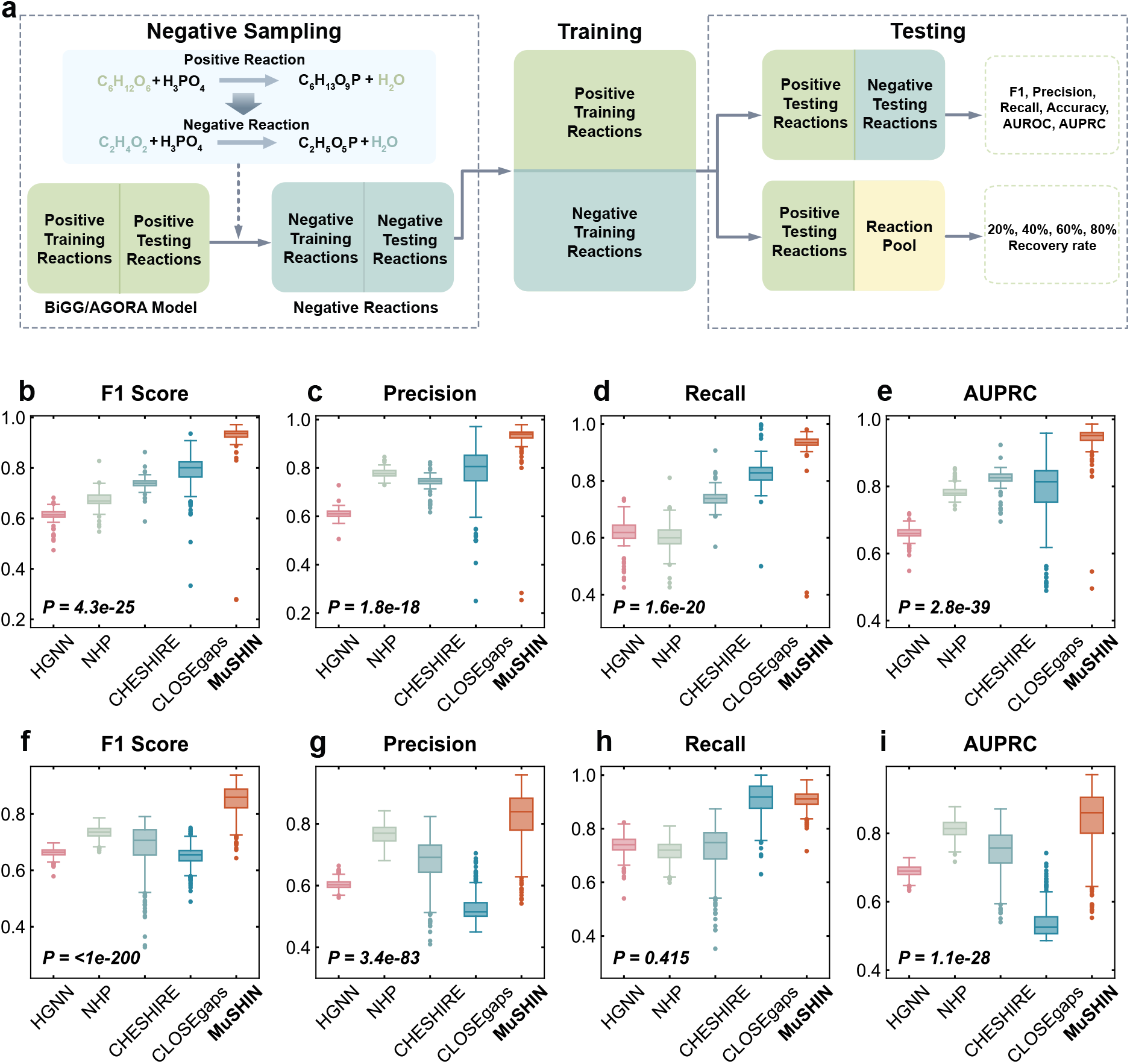
Overview and internal validation on BiGG and AGORA GEMs. **a**. Workflow of internal validation. Positive reactions from BiGG/AGORA GEMs are split into training and testing sets, while negative reactions are generated by perturbing valid reactions to create chemically invalid ones. Models are trained to distinguish positives from negatives and evaluated using statistical metrics. **b–e**. Internal validation results on 108 BiGG GEMs, evaluated using six metrics (F1 Score, Precision, Recall, AUPRC). Each dot represents one GEM, averaged over 10 Monte Carlo runs. MuSHIN consistently outperforms HGNN, NHP, CHESHIRE, and CLOSEgaps. **f–i**. Internal validation on 818 AGORA GEMs of gut bacteria using the same evaluation setup and metrics. Two-sided paired-sample t-tests were conducted between MuSHIN and the second-best baseline; exact p-values are reported. Each boxplot shows the median (central line), interquartile range (boxes), and variability across GEMs (whiskers and individual points). Source data are provided as a Source Data file.

On 108 GEMs from the BiGG database, MuSHIN exhibits consistent superiority across multiple evaluation metrics when benchmarked against four baseline methods (**Fig. 2b-e**). Specifically, MuSHIN achieves a median F1 score of 93.69%, precision of 93.98%, recall of 93.49%, and an AUPRC of 95.20%. Relative to the strongest baseline, CLOSEgaps, MuSHIN improves the F1 score by 17.01% (*P*-value = 4.3E-25; paired t-test), precision by 16.65% (*P*-value = 1.8E-18), recall by 12.86% (*P*-value = 1.6E-20), and AUPRC by 16.99% (*P* –value = 2.1E-27). These improvements reflect significant enhancements in both sensitivity (recall) and specificity (precision), critical for accurate gap-filling in GEM reconstruction. Performance margins over other baselines, such as CHESHIRE, NHP, and HGNN, are even more pronounced. Beyond its strong average performance, MuSHIN exhibits lower performance variance across GEMs of varying sizes, as indicated by tighter interquartile ranges, particularly in precision and AUPRC (**Fig. 2c**, e). This consistency suggests that MuSHIN is robust across networks of different complexity, in contrast to CLOSEgaps, which shows markedly higher variance, due to its sensitivity to network size and training instability in smaller models. Furthermore, MuSHIN maintains high performance under a range of evaluation settings, including different thresholding criteria, negative sampling strategies, and various sampling ratios, consistently surpassing all baselines (see Supplementary Note 3; Supplementary Figs. 1-5). Similar trends are observed on the 818 GEMs from the AGORA database, where MuSHIN again outperforms all baselines across all metrics (**Fig. 2f-i**). These results confirm that the advantages of MuSHIN are not confined to a specific dataset but instead generalize robustly across diverse reconstruction platforms, organismal systems, and metabolic network architectures, positioning MuSHIN as a state-of-the-art framework for accurate, scalable, and reliable reaction recovery in genome-scale metabolic modeling.

### MuSHIN enables robust recovery in highly incomplete GEMs

To assess the resilience of MuSHIN in scenarios of severe network incompleteness, we conducted a second internal validation experiment (**Fig. 2a**, bottom testing scheme). We selected nine representative GEMs from the BiGG database, each comprising approximately 2,000 reactions, and systematically removed 20%, 40%, 60%, or 80% of their reactions to simulate missing data. The remaining reactions were used for model training, supplemented with synthetically generated negative reactions at a 1:1 positive-to-negative ratio. For each level of degradation, we evaluated the model’s ability to recover missing reactions by measuring how many of the top *N* predicted reactions correctly matched to the *N* removed ones. MuSHIN was benchmarked against the same four baselines, CLOSEgaps, CHESHIRE, NHP, and HGNN, using identical model configurations and experimental protocols as in the first type of internal validation. This setup provides a stringent test of the model’s recoverability under varying degrees of sparsity.

MuSHIN consistently achieves high recovery rates across the selected GEMs and varying levels of missing rate, with an overall mean recovery approaching 90% (**Fig. 3**). When averaged across all conditions, MuSHIN outperforms the strongest baseline, CLOSEgaps, by 8.74%, demonstrating substantial and consistent gains over all competing methods. For example, in the model`STM_v1_0` (**Fig. 3h**), MuSHIN exceeds CLOSEgaps by roughly 10% across all tested removal rates (*P*-value = 1.3E-4). At the particularly challenging 60% removal level, MuSHIN maintains a recovery rate of 88.12%, whereas CLOSEgaps falls to 75.57%, underscoring the superior resilience of MuSHIN under extreme data sparsity. Across most GEMs and removal scenarios, the performance improvements of MuSHIN over baselines are statistically significant, with paired *P*-values below 0.001 in the majority of cases, and below 0.01 in the remainder. Additionally, the baseline methods exhibit notable variability in recovery performance as removal rates increase. In particular, HGNN fails to generalize under high levels of network incompleteness, often yielding recovery rates below 30% when 80% of reactions are withheld. In contrast, MuSHIN remains remarkably stable across all levels of incompleteness and model sizes, reflecting its capacity to generalize effectively even in data-scarce scenarios. These results highlight the robustness of MuSHIN in recovering complex metabolic structure from limited information and its suitability for reconstructing highly incomplete GEMs.

**Fig. 3:**
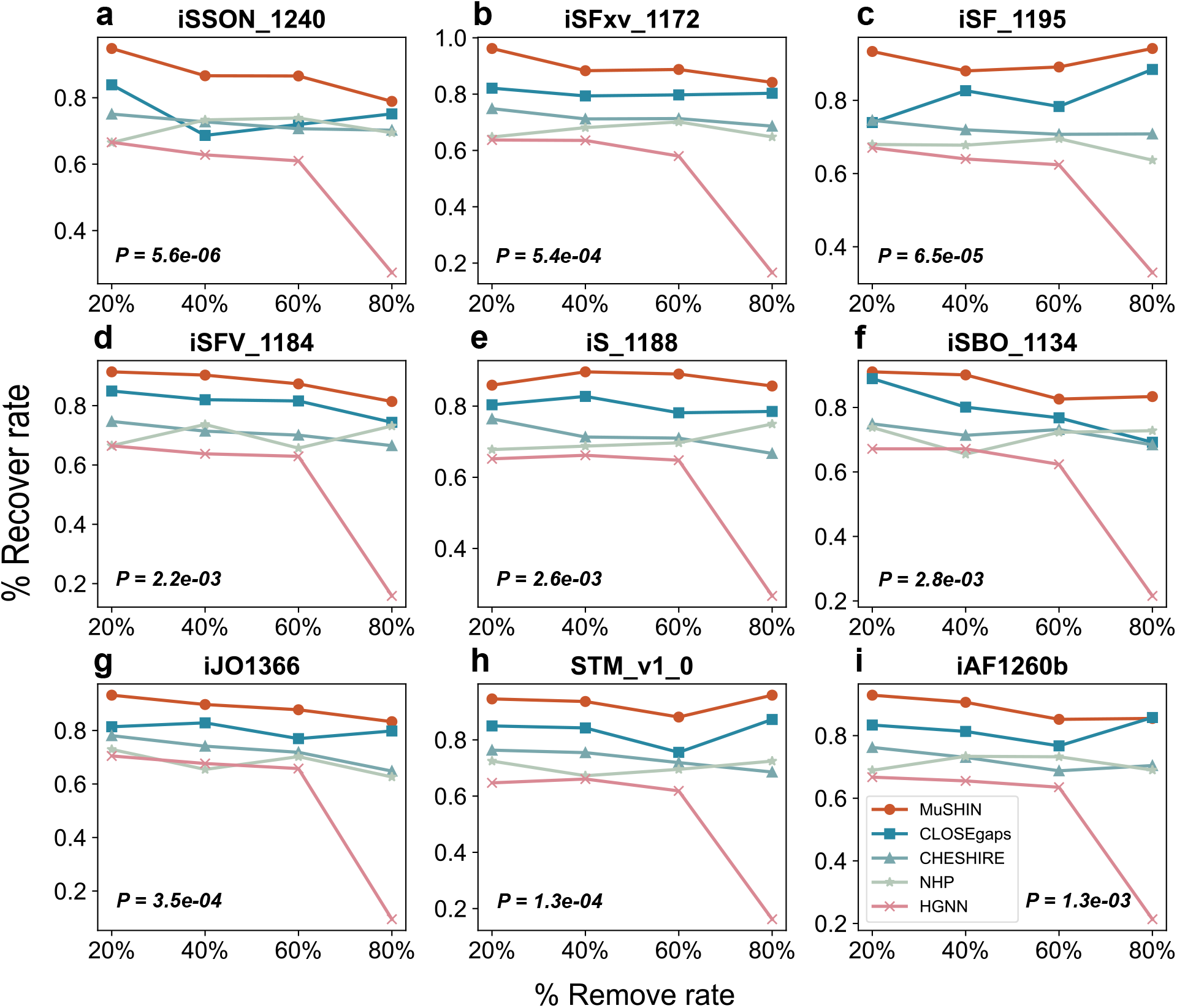
Recovery performance on highly incomplete GEMs. The recovery rates of MuSHIN and four baseline methods (HGNN, NHP, CHESHIRE, and CLOSEgaps) are evaluated across nine GEMs (`iAF1260b, iJO1366, iS_1188, iSBO_1134, iSF_1195, iSFV_1184, iSFxv_1172, iSSON_1240, STM_v1_0`) with varying levels of incompleteness (20%, 40%, 60%, and 80% of reactions removed). MuSHIN consistently demonstrates higher recovery rates compared to the baselines across all GEMs and removal rates, with particularly stable performance even under extreme scenarios (80% removal). Each data point represents the average recovery rate from 10 Monte Carlo simulations, while the lines illustrate trends across removal rates. Two-sided independent t-tests were performed between MuSHIN and CLOSEgaps for each model; exact p-values are shown.

### MuSHIN advances accurate phenotypic prediction in fermentation

To evaluate the biological relevance and practical utility of MuSHIN, we performed external validation to test whether gap-filling with MuSHIN enhances the prediction of actual metabolic phenotypes in draft GEMs against experimental data^12^. Our external validation workflow (**Fig. 4a**) begins by filtering candidate reactions from a universal reaction pool (e.g., the universal BiGG reaction pool), excluding those already present in the draft model or involving molecular oxygen. MuSHIN then scores the remaining candidates using a composite metric that integrates multiple evidence sources, including model-based scores and similarity coefficients, to prioritize reactions for gap-filling. We subsequently applied flux balance analysis to both the original draft and MuSHIN-augmented models to predict metabolic phenotypes. These predictions were rigorously compared to experimental phenotypic data using established classification metrics, enabling quantitative evaluation of MuSHIN’s contribution to improving metabolic phenotype prediction. Detailed steps, including similarity coefficients and flux balance analysis, can be found in Supplementary Note 5.

**Fig. 4:**
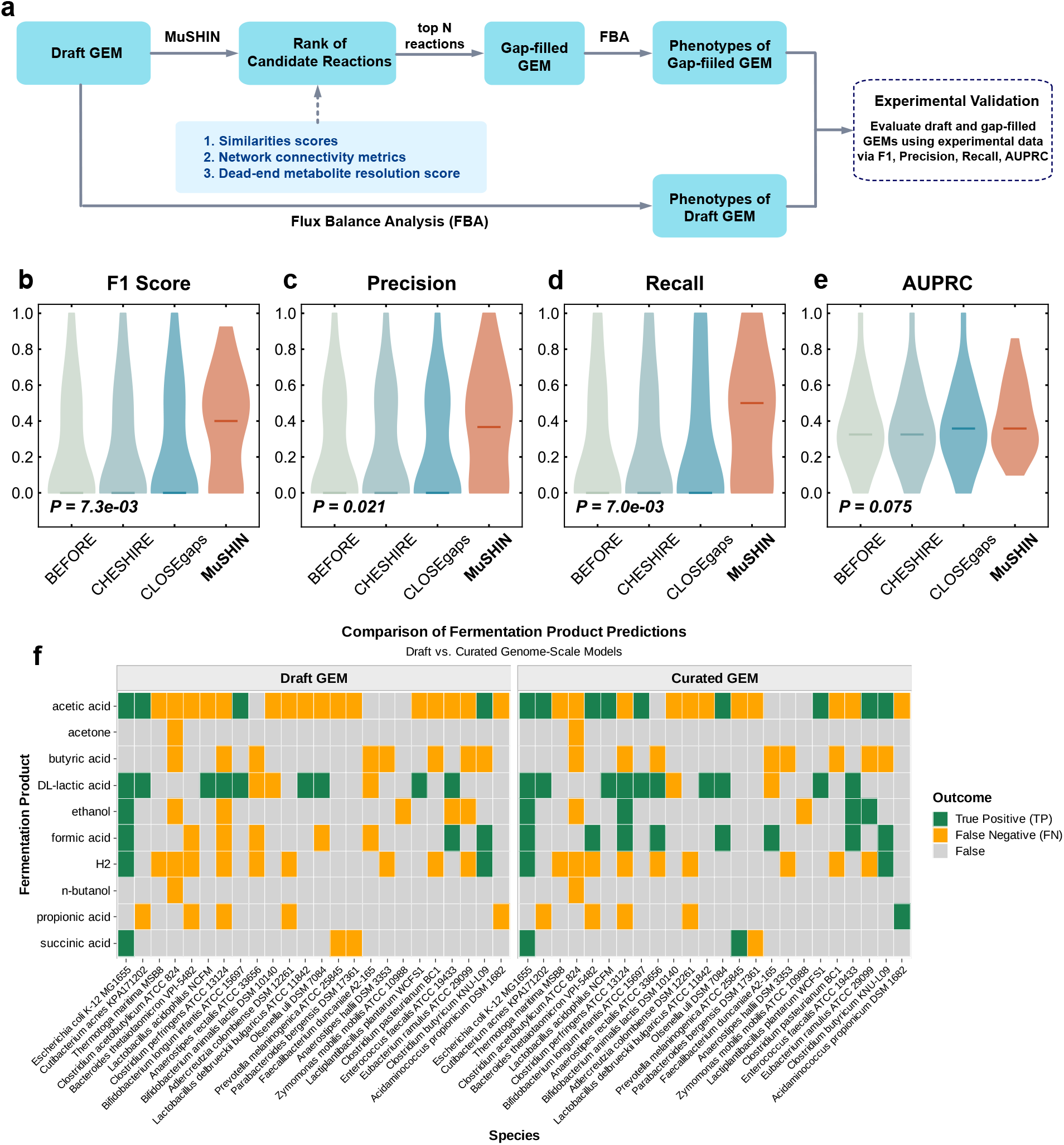
Overview and external validation performance. **a**. Workflow of external validation. MuSHIN ranks candidate reactions for draft GEMs, and top predictions are used to construct gap-filled GEMs. Flux balance analysis is applied to both draft and gap-filled models to simulate phenotypes, which are then compared with experimental data using classification metrics. **b–e**. External validation performance with 100 added reactions on anaerobic bacteria GEMs. Violin plots show F1 Score, Precision, Recall, and AUPRC for MuSHIN, CHESHIRE, CLOSEgaps, and the initial model (‘BEFORE’). MuSHIN outperforms CHESHIRE and the initial state, and is competitive or superior to CLOSEgaps across all metrics. Each point represents the mean result from 10 Monte Carlo simulations. Two-sided Wilcoxon signed-rank tests were conducted between MuSHIN and the second-best baseline method; exact p-values are shown in each panel. Each boxplot shows the median (central line), interquartile range (boxes), and variability across GEMs (whiskers and individual points). **f**. Validation of predicted fermentation products. Heatmap compares MuSHIN-predicted fermentation capabilities with experimentally known outcomes across 24 bacterial strains and 10 common fermentation products. Green cells indicate known products successfully recovered by MuSHIN, while orange boxes mark false-positive predictions introduced during gap-filling.

We applied MuSHIN to a set of 24 draft GEMs (Supplementary Table 1) derived from anaerobic bacterial species^41^ and evaluated its ability to improve phenotypic predictions of fermentation (see Supplementary Note 5). Specifically, we incorporated top 100 reactions prioritized by MuSHIN (MuSHIN-100) into each draft model and benchmarked the resulting performance against three comparators: the unmodified draft models (BEFORE), CHESHIRE-augmented models (CHESHIRE-100)^12^, and CLOSEgaps-augmented models (CLOSEgaps-^13^. As shown in **Fig. 4b-e**, MuSHIN-augmented models exhibit substantial improvements across multiple evaluation metrics, including F1 score, precision, recall, and AUPRC, compared to all the other models. MuSHIN achieves median values exceeding 40% for F1 score, precision, and recall, whereas the corresponding metrics for all other models remain zero (*P*-values = 7.3E-3, 2.1E-2, and 7.0E-3, respectively; Wilcoxon signed-rank test), underscoring the unique advantages conferred by MuSHIN in recovering biologically meaningful phenotypic predictions. Notably, previous studies have shown that both CHESHIRE and CLOSEgaps require the addition of at least 200 reactions to observe any significant improvement in predictive performance^12,13^. A detailed comparison of fermentation product predictions between draft and MuSHIN-augmented models is shown in **Fig. 4f**, where MuSHIN successfully corrects several false-negative predictions from the draft GEMs, enhancing agreement with experimentally observed fermentation phenotypes. Additionally, MuSHIN consistently outperforms CHESHIRE, CLOSEgaps, and the non-specific addition of 100 universal reactions (Supplementary Fig. 7). These findings highlight MuSHIN’s predictive strength and practical utility in improving the functional fidelity of GEMs under experimentally grounded and biologically realistic conditions, making it a potential tool for advancing biotechnology, microbial ecology, and systems biology research.

### MuSHIN resolves critical gaps in GEMs

To further understand the underlying biological advances provided by MuSHIN, we performed enrichment analyses for reactions prioritized by MuSHIN in the previous external experiment. Our results reveal a striking enrichment of transporter and exchange reactions. Among unique reactions proposed by MuSHIN (e.g., in the MuSHIN-100 set), 49.92% are annotated as transport or exchange, representing a significant enrichment compared to their prevalence in universal reaction databases (28.1% in the BiGG universal reaction pool; *P*-value *<* 1.0E-50; Fisher’s exact test). This selective enrichment aligns with the “membrane bottleneck hypothesis” in metabolic engineering, where transport limitations often constrain metabolic flux more than enzymatic capacity, indicating that MuSHIN selectively identifies boundary reactions in a non-random, biologically grounded manner.

To elucidate the mechanistic basis for MuSHIN’s improved phenotypic predictions, we compared its performance against the CHESHIRE-100 baseline across 24 selected genomes (Supplementary Table 1). MuSHIN improves the phenotypic prediction F1 score in 11 of these genomes, yielding substantial gains in forecasting specific metabolic products. Notably, MuSHIN enables the correct prediction of formic acid in five genomes and acetic acid in five other genomes where CHESHIRE-100 fails. Additional significant improvements are observed for ethanol, which is uniquely predicted by MuSHIN in three distinct genomes missed by CHESHIRE. Furthermore, DL-lactic acid, succinic acid, and propionic acid are each correctly identified in one additional genome. Collectively, MuSHIN introduces 16 new correct product predictions absent in the CHESHIRE-100 results, highlighting its superior capacity to recover missing metabolic functions. Further analysis (Supplementary Table 2) reveals that these improvements are likely explained by MuSHIN’s addition of specific transporter families, which enhance substrate secretion or cofactor availability. Key examples include transporters for formate (FORt2^∗^, FORt3^∗^), CO_2_*/*O_2_ exchange (CO2tex, O2tex), Zn^2+^ (ZN2t^∗^, Zn2tex), ethanol (ETOHt^∗^), and fumarate/malate (FUMt^∗^, MALt^∗^). For instance, improved formic acid secretion is largely due to the addition of specific formate transporters (FORt2^∗^, FORt3^∗^) in six genomes^42^, often coupled with CO_2_ exchange transporters (CO2tex) that facilitate carbon metabolism^43^. Similarly, enhanced ethanol production is led by the addition of ethanol-specific transporters (ETOHt2r^∗^, ETOHt3^∗^) in five genomes, enabling efficient product efflux^44^.

Beyond single-transporter corrections, MuSHIN uniquely resolves complex metabolic gaps. A notable instance in *GCF 000144405*.*1* involves the restoration of succinate production through coordinated addition of fumarate transporters (FUMt2 3pp, FUMt2 2pp, FUMtpp), facilitating the reductive branch of the tricarboxylic acid cycle (TCA) under anaerobic conditions^45^. Similarly, in *GCF 000013285*.*1*, improvements in both ethanol and formic acid production are achieved through the addition of ethanol-specific transporters (ETOHt2rpp, ETOHt3, ETOHt2r) and formate transporters (FORt2pp, FORt3, FORt2)^46^, demonstrating MuSHIN’s ability to enhance multiple fermentation pathways simultaneously. In genomes like *GCF 000392875*.*1* and *GCF 000469345*.*1*, MuSHIN enhances ethanol production even without the addition of dedicated ethanol transporters, suggesting the involvement of alternative transport mechanisms or metabolic rerouting. This synergy highlights MuSHIN’s strength in uncovering both direct transporter-mediated solutions and indirect metabolic enhancements–complex scenarios that simpler gap-filling methods often miss. Notably, zinc transporters (ZN2t^∗^, Zn2tex) are added across all eleven genomes, indicating their fundamental role as cofactor suppliers for enhanced enzymatic activity in metabolic pathway restoration. The universal requirement for zinc transporters reflects the metal’s role as a cofactor in over 300 enzymes, particularly those involved in central carbon metabolism and redox reactions^45^.

In summary, these findings demonstrate that MuSHIN’s superior performance arises from addressing a fundamental limitation in current gap-filling approaches: the systematic underestimation of membrane transport constraints. By prioritizing functionally critical transport and exchange reactions that alleviate these bottlenecks, and resolving complex multi-reaction gaps through coordinated transporter addition, MuSHIN reinstates complete metabolic branches while maintaining the stoichiometric and thermodynamic constraints essential for biological fidelity.

## Discussion

In this work, we introduced MuSHIN (**Mu**lti-way **S**MILES-based **H**ypergraph **I**nterface **N**etwork), a deep hypergraph learning framework that integrates metabolic network topology with biochemical domain knowledge to address the persistent challenge of reaction incompleteness in GEMs. By representing metabolic networks as hypergraphs, MuSHIN captures the intrinsic high-order relationships among metabolites and reactions, offering a more faithful abstraction of biochemical systems than traditional pairwise graph representations. In addition, MuSHIN incorporates chemically informed molecular embeddings derived from SMILES strings, enabling the model to reason about molecular structure and reactivity. A dynamic attention mechanism further enhances this process by selectively emphasizing the most informative topological and biochemical features, allowing MuSHIN to adaptively prioritize relevant signals during training and prediction. The contributions of these components are further supported by ablation results (Supplementary Fig. 6). Compared to recent state-of-the-art methods such as CHESHIRE and CLOSEgaps, which primarily rely on reaction co-occurrence patterns or curated reaction similarity metrics, MuSHIN offers a fundamentally different learning paradigm, which allows MuSHIN to generalize more effectively, particularly in sparse or incompletely annotated networks, and establishes a robust foundation for scalable and biologically grounded metabolic network reconstruction.

The effectiveness of MuSHIN was rigorously evaluated through both internal and external validation. Internally, we conducted extensive benchmarking on a large collection of 926 high- and intermediate-quality GEMs, where reactions were systematically and artificially removed to simulate real-world incompleteness. Across multiple quantitative metrics, MuSHIN consistently outperforms existing state-of-the-art hypergraph-based gap-filling methods such as CHESHIRE and CLOSEgaps. Notably, MuSHIN maintains strong predictive performance even under conditions of extreme network sparsity, demonstrating its ability to infer missing reactions in challenging scenarios where conventional approaches falter. Externally, we applied MuSHIN to 24 draft GEMs focused on microbial fermentation pathways and validated the predicted reaction additions by comparing model-derived phenotypic outputs with independent experimental measurements. Incorporation of 100 reactions predicted by MuSHIN leads to significant improvements in fermentation phenotype predictions, whereas other methods such as CHESHIRE and CLOSEgaps fail to produce noticeable enhancements. This demonstrates MuSHIN’s superior ability to identify functionally relevant reactions that directly translate to more accurate and biologically meaningful phenotypic outcomes.

Enhancing the completeness and fidelity of GEMs holds transformative potential across diverse domains of biology, biotechnology, and medicine. In biotechnology, high-quality GEMs enable rational strain engineering by revealing metabolic bottlenecks and guiding the optimization of biosynthetic pathways for the production of pharmaceuticals, biofuels, and other high-value compounds. In the context of human health, GEMs serve as powerful tools for precision medicine, allowing the modeling of tissue- or patient-specific metabolism to uncover disease mechanisms, identify biomarkers, and inform therapeutic interventions. In microbial ecology and synthetic biology, more comprehensive metabolic reconstructions facilitate the design of synthetic consortia and enable the strategic allocation of metabolic functions across community members. By advancing automated, data-driven approaches for metabolic gap-filling such as MuSHIN, we not only improve the structural and functional accuracy of GEMs but also unlock their full potential as predictive frameworks for understanding and engineering complex biological systems.

Building on MuSHIN’s strong performance, several avenues offer promising opportunities for further development. First, while the framework effectively integrates molecular structure and metabolic network topology, it currently lacks integration of additional biological layers, such as gene-protein-reaction associations and regulatory interactions, which could be harnessed to construct more comprehensive, multi-modal knowledge hypergraphs. Second, although SMILES-based embeddings capture chemical properties, they may not fully reflect enzyme-specific context or regulatory constraints, which are critical for accurate functional interpretation. Third, MuSHIN currently treats gap-filling as a static prediction task and does not incorporate dynamic cellular states or context-dependent metabolic shifts, which are increasingly important for modeling complex biological systems. Pursuing these directions will support broader applicability of MuSHIN and further advance the automation and accuracy of GEM reconstruction.

## Methods

### Hypergraphs

Hypergraphs extend traditional graphs by incorporating hyperedges (also known as hyperlinks) that can connect any number of nodes^30^. This flexibility allows hypergraphs to model complex correlations in real-world data that go beyond simple pairwise interactions^37^. Formally, an unweighted hypergraph ℋ = {𝒱, ℰ} comprises a set of nodes 𝒱 = {*v*_1_, *v*_2_, …, *v*_*n*_} and a set of hyperedges ℰ = {*e*_1_, *e*_2_, …, *e*_*m*_}, where each hyperedge *e*_*p*_ ⊆ 𝒱 for *p* = 1, 2, …, *m*. Two nodes are considered adjacent if they share a common hyperedge, and a hypergraph is connected if there exists a path between any two nodes via a sequence of hyperedges. The incidence matrix of a hypergraph, denoted by **H** ∈ {0, 1}^*n*×*m*^, captures the relationships between nodes and hyperedges. Specifically, the entry **H**_*ip*_ is set to one if node *v*_*i*_ is part of hyperedge *e*_*p*_, and zero otherwise.

### Negative reaction generation

Predicting missing reactions in metabolic networks necessitates a dataset containing both positive (existing) and negative (non-existing) reactions. We developed a negative sampling strategy that generates invalid reactions by perturbing known reactions. Given a set of known reactions ℛ^+^ = {*r*_1_, *r*_2_, …, *r*_*N*_ }, where each reaction *r*_*i*_ comprises specific reactants and products, we generate negative reactions through the following steps: (i) aggregate a metabolite pool ℳ by combining metabolites from ℛ^+^ with those from external databases; (ii) for each reaction *r*_*i*_ ∈ ℛ^+^, randomly select a fraction *α* of its metabolites for replacement; (iii) substitute the selected metabolites with randomly chosen ones from ℳ \ *r*_*i*_ (i.e., metabolites not present in *r*_*i*_); (iv) ensure that the newly generated reaction 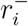 does not exist in ℛ^+^.

This procedure yields a set of negative reactions 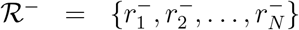 that are structurally similar to positive reactions but are chemically invalid due to improbable metabolite combinations^47, 48^. Each reaction is represented by a stoichiometric vector **s**_*i*_ ∈ ℝ^*M*^, where *M* is the total number of unique metabolites, and each component *s*_*i,j*_ represents the stoichiometric coefficient *n*_*j*_ of metabolite *j*:

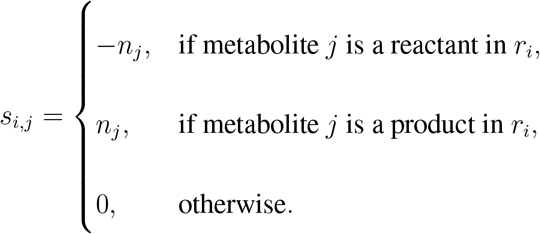

The complete dataset is ℛ = ℛ^+^ ∪ ℛ^−^, with each reaction labeled as follows:

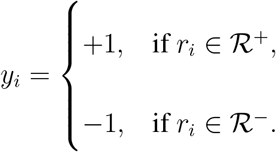

### Reaction embedding using RXNFP

RXNFP^37^ is a transformer-based model pre-trained on chemical reaction data, designed to encode reaction SMILES strings into numerical representations that capture underlying chemical transformations and mechanistic patterns. Each reaction SMILES string is tokenized and processed through multiple transformer layers, producing a fixed-length reaction fingerprint (typically 256-dimensional) suitable for downstream tasks such as reaction classification and pathway prediction. Mathematically, the RXNFP model processes a tokenized reaction SMILES sequence *r* = [*r*_1_, *r*_2_, …, *r*_*N*_] through transformer layers as follows:

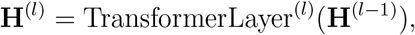

where **H**^(0)^ represents the initial token embeddings, and **X**^(*l*)^ ∈ ℝ^*N* ×*d*^ denotes the hidden states at layer *l*, with *d* being the hidden size. The final reaction fingerprint **f** ∈ ℝ^*d*^ is obtained by applying a pooling operation over the last layer’s hidden states **f** = Pooling(**H**^(*L*)^). By integrating RXNFP-based embeddings, MuSHIN effectively incorporates chemical reaction patterns into the hypergraph model, improving its ability to predict missing reactions.

### Metabolite embedding using ChemBERTa

Similarly, ChemBERTa^38^ is a transformer-based language model pre-trained on molecular data, which encodes metabolite SMILES strings into meaningful vector representations. ChemBERTa captures complex structural and chemical relationships, providing robust numerical features for metabolic network analysis. Metabolite data is sourced from the BiGG database^10^, and corresponding SMILES representations are retrieved via ChEBI IDs^47^. If unavailable, external databases such as KEGG are consulted. Each SMILES string is tokenized and processed through multiple transformer layers, producing a fixed-length (768-dimensional) vector representation. Mathematically, for a tokenized SMILES sequence *s* = [*s*_1_, *s*_2_, …, *s*_*N*_], the ChemBERTa model generates hidden states at each transformer layer. The hidden states from the final layer *L* are denoted as

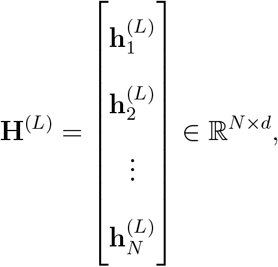

where 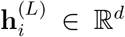 represents the hidden state of the *i*th token, and *d* = 768. The metabolite’s vector representation **v** ∈ ℝ^*d*^ is computed by averaging these hidden states 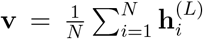.

By leveraging ChemBERTa-based embeddings, MuSHIN incorporates rich molecular properties into its hypergraph representation, allowing for a more accurate characterization of metabolite relationships within metabolic networks. The combination of RXNFP and ChemBERTa ensures that MuSHIN effectively captures both topological and chemical features, enabling superior performance in predicting missing reactions.

### Hypergraph neural network architecture

Various hypergraph neural network models have been proposed^27, 33, 49–52^. However, most of these approaches simplify the problem by decomposing hyperedges into pairwise interactions and then applying standard graph neural networks, which can lead to a loss of high-order structural information. To effectively model the complex interactions within the metabolic network, we propose a novel hypergraph neural network with a dual attention mechanism that leverages both node and hyperedge features. Our architecture integrates advanced attention mechanisms to capture higher-order dependencies and interactions within the hypergraph. Specifically, the model iteratively refines feature representations through a combination of Node-to-Edge and Edge-to-Node attention mechanisms, allowing MuSHIN to dynamically learn the relationships between metabolites and reactions.

The Node-to-Edge attention mechanism aggregates information from nodes to hyperedges, updating hyperedge features based on the features of their incident nodes. This enables hyperedges to focus on the most relevant information from connected nodes, weighted by learned attention coefficients. Given node features 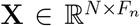 and the incidence matrix **H** ∈ {0, 1}^*N* ×*M*^, where *N* is the number of nodes, *M* is the number of hyperedges, and *F*_*n*_ is the dimensionality of node features, we first transform the node features into a shared latent space

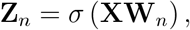

where 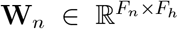 is a learnable weight matrix, *F*_*h*_ is the hidden dimension, and *σ* is an activation function (e.g., ReLU). For each hyperedge *e*, we compute attention coefficients for its incident nodes

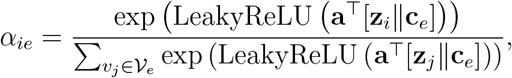

where **z**_*i*_ is the transformed feature of node 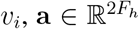 is a learnable attention vector, ∥ denotes vector concatenation, 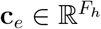 is a learnable hyperedge representation (initialized randomly or as zeros), and 𝒱_*e*_ denotes the set of nodes connected to hyperedge *e*. The hyperedge feature **h**_*e*_ is then updated by aggregating the node features weighted by the attention coefficients 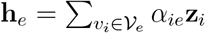. This mechanism allows hyperedges to emphasize the most informative nodes, enhancing their ability to represent complex relationships within the hypergraph.

The Edge-to-Node attention mechanism updates node features by aggregating information from the hyperedges they are connected to. By attending to the most relevant hyperedges, nodes can incorporate higher-order relational information into their representations. First, we transform both node and hyperedge features into a shared latent space

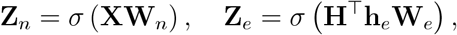

where 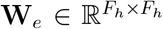 is a learnable weight matrix for hyperedges. For each node *i*, the attention coefficients for connected hyperedges are computed as

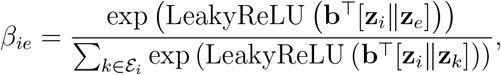

where **z**_*e*_ is the transformed feature of hyperedge 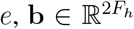 is a learnable attention vector, and ℰ_*i*_ denotes the set of hyperedges connected to node *i*. The updated node feature 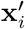 is obtained by aggregating the hyperedge features weighted by the attention coefficients and adding a residual connection 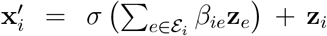. This approach allows nodes to integrate contextual information from relevant hyperedges, resulting in richer feature representations.

We iteratively apply the Node-to-Edge and Edge-to-Node attention mechanisms to progressively refine both node and hyperedge features. At each iteration, updated node features inform the hyperedge features, and updated hyperedge features subsequently enhance the node features. This iterative process facilitates deep integration of structural information across the hypergraph, capturing complex dependencies and interactions. The iterative refinement process is formalized as follows:

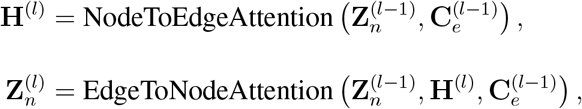

for *l* = 1, 2, …, *L*, where 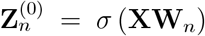 are the initial node features, **H**^(*l*)^ represents the hyperedge features at layer *l*, and 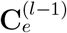 are the hyperedge representations from the previous layer. This iterative refinement enhances the model’s ability to capture higher-order dependencies and interactions within the hypergraph, leading to improved performance on downstream tasks such as reaction prediction and network analysis (see Supplementary Note 3 for a detailed prediction workflow).

## Data availability

The datasets used and analyzed during the current study are included within this article and its supplementary information files. The raw data were collected from publicly available databases: ChEBI (https://www.ebi.ac.uk/chebi/), BiGG Models (http://bigg.ucsd.edu/), AGORA Models (https://www.vmh.life). More details can be found in Supplementary Note 5.

## Code availability

The source code for our framework is available at Github [https://github.com/cyixiao/MuSHIN].

## References

1. Ines Thiele, Nathan D Price, Thuy D Vo, and Bernhard Ø Palsson. Candidate metabolic network states in human mitochondria. FEBS Letters, 579(30):6435–6441, 2005.

2. Ines Thiele, Neema Jamshidi, Ronan MT Fleming, and Bernhard Ø Palsson. Genome-scale reconstruction of escherichia coli’s transcriptional and translational machinery: A knowledge base, its mathematical formulation, and its functional characterization. PLoS Computational Biology, 5(3):e1000312, 2009.

3. Sang Yup Lee and Hyun Uk Kim. Systems strategies for developing industrial microbial strains. Nature Biotechnology, 33(10):1061–1072, Oct 2015.

4. Changdai Gu, Gi Bae Kim, Won Jun Kim, Hyun Uk Kim, and Sang Yup Lee. Current status and applications of genome-scale metabolic models. Genome biology, 20:1–18, 2019.

5. Christian Lieven, Moritz E Beber, Brett G Olivier, Frank T Bergmann, Meric Ataman, Parizad Babaei, Jennifer A Bartell, Lars M Blank, Siddharth Chauhan, Kevin Correia, et al. Memote for standardized genome-scale metabolic model testing. Nature biotechnology, 38(3):272–276, 2020.

6. Evangelos Simeonidis and Nathan D Price. Genome-scale modeling for metabolic engineering. Journal of Industrial Microbiology and Biotechnology, 42(3):327–338, 2015.

7. Byoungjin Kim, Won Jun Kim, Dong In Kim, and Sang Yup Lee. Applications of genome-scale metabolic network model in metabolic engineering. Journal of industrial microbiology and biotechnology, 42(3):339–348, 2015.

8. Vytautas Raškevičius, Valeryia Mikalayeva, Ieva Antanavičiūtė, Ieva Ceslevičienė, Vytenis Arvydas Skeberdis, Visvaldas Kairys, and Sergio Bordel. Genome scale metabolic models as tools for drug design and personalized medicine. PloS one, 13(1):e0190636, 2018.

9. Jonathan L Robinson and Jens Nielsen. Anticancer drug discovery through genome-scale metabolic modeling. Current Opinion in Systems Biology, 4:1–8, 2017.

10. Zachary A King, Justin Lu, Andreas Dräger, Philip Miller, Stephen Federowicz, Joshua A Lerman, Ali Ebrahim, Bernhard O Palsson, and Nathan E Lewis. Bigg models: A platform for integrating, standardizing and sharing genome-scale models. Nucleic acids research, 44(D1):D515–D522, 2016.

11. Jens Nielsen and Jay D. Keasling. Engineering cellular metabolism. Cell, 164(6):1185–1197, 2016.

12. Can Chen, Chen Liao, and Yang-Yu Liu. Teasing out missing reactions in genome-scale metabolic networks through hypergraph learning. Nature Communications, 14(1):2375, 2023.

13. Xiaoyi Liu, Hongpeng Yang, Chengwei Ai, Ruihan Dong, Yijie Ding, Qianqian Yuan, Jijun Tang, and Fei Guo. A generalizable framework for unlocking missing reactions in genome-scale metabolic networks using deep learning. arXiv preprint arXiv:2409.13259, 2024.

14. Shu Pan and Jennifer L Reed. Advances in gap-filling genome-scale metabolic models and model-driven experiments lead to novel metabolic discoveries. Current opinion in biotechnology, 51:103–108, 2018.

15. Matthew N Benedict, Michael B Mundy, Christopher S Henry, Nicholas Chia, and Nathan D Price. Likelihood-based gene annotations for gap filling and quality assessment in genome-scale metabolic models. PLoS computational biology, 10(10):e1003882, 2014.

16. Peter D Karp, Daniel Weaver, and Mario Latendresse. How accurate is automated gap filling of metabolic models? BMC systems biology, 12:1–11, 2018.

17. Daniel Machado, Sergej Andrejev, Michele Tramontano, and Kiran R Patil. Fast automated reconstruction of genome-scale metabolic models for microbial species and communities. Nucleic acids research, 46(15):7542–7553, 2018.

18. Changdai Gu, Gi Bae Kim, Won Jun Kim, Hyun Uk Kim, and Sang Yup Lee. Current status and applications of genome-scale metabolic models. Genome Biology, 20(1):121, June 2019.

19. Jeffrey D Orth and Bernhard Ø Palsson. Systematizing the generation of missing metabolic knowledge. Biotechnology and bioengineering, 107(3):403–412, 2010.

20. Wheaton L Schroeder and Rajib Saha. Optfill: a tool for infeasible cycle-free gapfilling of stoichiometric metabolic models. IScience, 23(1), 2020.

21. Sylvain Prigent, Clémence Frioux, Simon M Dittami, Sven Thiele, Abdelhalim Larhlimi, Guillaume Collet, Fabien Gutknecht, Jeanne Got, Damien Eveillard, Jérémie Bourdon, et al. Meneco, a topology-based gap-filling tool applicable to degraded genome-wide metabolic networks. PLoS computational biology, 13(1):e1005276, 2017.

22. Vinay Satish Kumar, Madhukar S Dasika, and Costas D Maranas. Optimization based automated curation of metabolic reconstructions. BMC bioinformatics, 8:1–16, 2007.

23. Christopher S Henry, Matthew DeJongh, Aaron A Best, Paul M Frybarger, Ben Linsay, and Rick L Stevens. High-throughput generation, optimization and analysis of genome-scale metabolic models. Nature biotechnology, 28(9):977–982, 2010.

24. Alexandre Almeida, Alex L Mitchell, Miguel Boland, Samuel C Forster, Gregory B Gloor, Aleksandra Tarkowska, Trevor D Lawley, and Robert D Finn. A new genomic blueprint of the human gut microbiota. Nature, 568(7753):499–504, 2019.

25. Can Chen and Yang-Yu Liu. A survey on hyperlink prediction. IEEE Transactions on Neural Networks and Learning Systems, 35(11):15034–15050, 2024.

26. Yifan Feng, Haoxiang You, Zhirong Zhang, Rongrong Ji, and Yue Gao. Hypergraph neural networks. In Proceedings of the AAAI Conference on Artificial Intelligence, volume 33, pages 3558–3565, 2019.

27. Yiwei Bai, Hao Ding, Yifan Sun, Wenyu Wang, and Yihong Gong. Hypergraph convolution and hypergraph attention. Pattern Recognition, 110:107637, 2021.

28. Can Chen, Amit Surana, Anthony M Bloch, and Indika Rajapakse. Controllability of hypergraphs. IEEE Transactions on Network Science and Engineering, 8(2):1646–1657, 2021.

29. Can Chen and Indika Rajapakse. Tensor entropy for uniform hypergraphs. IEEE Transactions on Network Science and Engineering, 7(4):2889–2900, 2020.

30. Claude Berge. Hypergraphs: combinatorics of finite sets, volume 45. Elsevier, 1984.

31. Dengyong Zhou, Jiayuan Huang, and Bernhard Schölkopf. Learning with hypergraphs: Clustering, classification, and embedding. Advances in neural information processing systems, 19, 2006.

32. Yue Gao, Zizhao Zhang, Haojie Lin, Xibin Zhao, Shaoyi Du, and Changqing Zou. Hypergraph learning: Methods and practices. IEEE Transactions on Pattern Analysis and Machine Intelligence, 44(5):2548–2566, 2020.

33. Yifan Feng, Haoxuan You, Zizhao Zhang, Rongrong Ji, and Yue Gao. Hypergraph neural networks, 2019.

34. Muhan Zhang, Zhicheng Cui, Shali Jiang, and Yixin Chen. Beyond link prediction: Predicting hyperlinks in adjacency space. In Proceedings of the AAAI conference on artificial intelligence, volume 32, 2018.

35. Govind Sharma, Prasanna Patil, and M Narasimha Murty. C3mm: Clique-closure based hyperlink prediction. In IJCAI, volume 20, pages 3364–3370, 2020.

36. Naganand Yadati, Vikram Nitin, Madhav Nimishakavi, Prateek Yadav, Anand Louis, and Partha Talukdar. Nhp: Neural hypergraph link prediction. In Proceedings of the 29th ACM international conference on information & knowledge management, pages 1705–1714, 2020.

37. Philippe Schwaller, Daniel Probst, Alain C Vaucher, Vishnu H Nair, David Kreutter, Teodoro Laino, and Jean-Louis Reymond. Mapping the space of chemical reactions using attention-based neural networks. Nature machine intelligence, 3(2):144–152, 2021.

38. Seyone Chithrananda, Gabriel Grand, and Bharath Ramsundar. Chemberta: large-scale self-supervised pretraining for molecular property prediction. arXiv preprint arXiv:2010.09885, 2020.

39. A Vaswani. Attention is all you need. Advances in Neural Information Processing Systems, 2017.

40. Stefanía Magnúsdóttir, Almut Heinken, Laura Kutt, Dmitry A Ravcheev, Eugen Bauer, Alberto Noronha, Kacy Greenhalgh, Christian Jäger, Joanna Baginska, Paul Wilmes, et al. Generation of genome-scale metabolic reconstructions for 773 members of the human gut microbiota. Nature biotechnology, 35(1):81–89, 2017.

41. David B Bernstein, Snorre Sulheim, Eivind Almaas, and Daniel Segrè. Addressing uncertainty in genome-scale metabolic model reconstruction and analysis. Genome Biology, 22:1–22, 2021.

42. Wei Lü, Juan Du, Nikola J Schwarzer, Elke Gerbig-Smentek, Oliver Einsle, and Susana LA Andrade. The formate channel foca exports the products of mixed-acid fermentation. Proceedings of the National Academy of Sciences, 109(33):13254–13259, 2012.

43. Max van ‘t Hof, Omkar S Mohite, Jonathan M Monk, Tilmann Weber, Bernhard O Palsson, and Morten OA Sommer. High-quality genome-scale metabolic network reconstruction of probiotic bacterium escherichia coli nissle 1917. BMC bioinformatics, 23(1):566, 2022.

44. Xiao Bu, Jing-Yuan Lin, Jing Cheng, Dong Yang, Chang-Qing Duan, Mattheos Koffas, and Guo-Liang Yan. Engineering endogenous abc transporter with improving atp supply and membrane flexibility enhances the secretion of β-carotene in saccharomyces cerevisiae. Biotechnology for Biofuels, 13:1–14, 2020.

45. Antoine Danchin. Zinc, an unexpected integrator of metabolism? Microbial Biotechnology, 13(4):895–898, 2020.

46. Wei Lü, Juan Du, Nikola J. Schwarzer, Elke Gerbig-Smentek, Oliver Einsle, and A. Andrade, Susana LA Andrade. The formate channel foca exports the products of mixed-acid fermentation. Proceedings of the National Academy of Sciences, 109(33):13254–13259, 2012.

47. Janna Hastings, Paula De Matos, Adriano Dekker, Marcus Ennis, Bhavana Harsha, Namrata Kale, Venkatesh Muthukrishnan, Gareth Owen, Steve Turner, Mark Williams, et al. The chebi reference database and ontology for biologically relevant chemistry: enhancements for 2013. Nucleic acids research, 41(D1):D456–D463, 2012.

48. Tomas Mikolov. Efficient estimation of word representations in vector space. arXiv preprint arXiv:1301.3781, 2013.

49. Yue Gao, Yifan Feng, Shuyi Ji, and Rongrong Ji. Hgnn+: General hypergraph neural networks. IEEE Transactions on Pattern Analysis and Machine Intelligence, 45(3):3181–3199, 2022.

50. Jianwen Jiang, Yuxuan Wei, Yifan Feng, Jingxuan Cao, and Yue Gao. Dynamic hypergraph neural networks. In IJCAI, pages 2635–2641, 2019.

51. Sunwoo Kim, Soo Yong Lee, Yue Gao, Alessia Antelmi, Mirko Polato, and Kijung Shin. A survey on hypergraph neural networks: An in-depth and step-by-step guide. In Proceedings of the 30th ACM SIGKDD Conference on Knowledge Discovery and Data Mining, pages 6534–6544, 2024.

52. Jaehyuk Yi and Jinkyoo Park. Hypergraph convolutional recurrent neural network. In Proceedings of the 26th ACM SIGKDD international conference on knowledge discovery & data mining, pages 3366–3376, 2020.

