## Supplemental material for "MuSHIN: A multi-way SMILES-based hypergraph inference network for metabolic model reconstruction"

### Supplementary Note 1: Existing methods of predicting missing reactions

**GapFind and GapFill**[1,2] are optimization-based approaches designed to address gaps in genome-scale metabolic models (GEMs). The GapFind algorithm identifies metabolites within the model that cannot be synthesized under any possible uptake conditions, thus exposing disconnected components in the network. To resolve these issues, GapFill leverages a curated database of reactions from multiple organisms and employs various strategies to restore connectivity. These include reversing the directionality of existing reactions, incorporating reactions from other organisms to compensate for missing functionalities, introducing external transport reactions to enable the uptake of inaccessible metabolites, and adding intracellular transport mechanisms to facilitate flow within multi-compartment models. Detailed descriptions of the underlying optimization formulations are available in the relevant literature.

**FastGapFill**[3] extends the functionality of GapFill and is the first scalable approach specifically designed for detecting and addressing gaps in compartmentalized GEMs. It begins by employing the FastCore algorithm to identify a near-minimal set of reactions required to ensure flux consistency in the input GEM. Following this, FastGapFill expands the compartmentalized model—defined as the version without any blocked reactions—by integrating reactions from a universal metabolic database, such as the Kyoto Encyclopedia of Genes and Genomes (KEGG) [4]. Finally, the method constructs a compact, flux-consistent subnetwork from the expanded global model, resulting in a finalized gap-filled model.

**Matrix boost algorithm (BoostGapFill)**[5] introduces a novel approach for gap filling by performing inference simultaneously in both the incidence and adjacency spaces of hypergraphs. It begins by approximating the incomplete adjacency matrix of a hypergraph using a combination of observed entries and iteratively estimated missing entries. The optimization process minimizes the difference between the predicted and observed adjacency matrices while applying a regularization term to ensure robustness. To further refine the selection of candidate hyperlinks from a predefined set, the method incorporates a matching step by solving an optimization problem that balances the reconstruction error of the adjacency matrix with the assignment of hyperlinks, which can be relaxed for computational efficiency Despite its strength in leveraging matrix factorization techniques, BoostGapFill exhibits scalability limitations. The requirement of a predefined candidate hyperlink set during training makes the method computationally impractical for large-scale databases such as Biochemical, Genetic and Genomic knowledge base (BiGG) [6]. Moreover, it lacks the flexibility to handle unseen hyperlinks during testing, limiting its applicability in scenarios that demand generalization beyond the training candidate pool.

**Coordinated Matrix Minimization (CMM)**[7] improves upon BoostGapFill by introducing a latent factor matrix, simplifying the algorithm and enhancing performance. CMM combines non-negative matrix factorization and least-squares matching to efficiently infer candidate hyperlinks that best complete the hypergraph. By representing the adjacency matrix as a product of latent factors, the optimization problem minimizes reconstruction error and identifies missing hyperlinks. This is achieved through an Expectation-Maximization framework, where constraints are relaxed for computational efficiency using standard optimization tools. Despite its advantages, CMM retains limitations in scalability and fails to generalize to unseen hyperlinks.

**Clique Closure-based Coordinated Matrix Minimization (C3MM)**[8] further extends CMM by leveraging clique-closure properties of hypergraphs. This approach incorporates additional structural information, enabling the identification of hyperlinks overlooked by CMM. The method solves an alternating optimization problem that incorporates clique-based regularization and minimizes reconstruction error while capturing common-neighbor information. C3MM shows strong performance in hyperlink prediction tasks but shares scalability and generalization limitations with its predecessors, indicating the need for more advanced techniques, such as deep learning, to address these issues.

**Node2Vec-mean (NVM)**[9] is a straightforward baseline for hyperlink prediction, utilizing a simple yet effective design. Starting with an incomplete hypergraph $\mathcal{H}$ containing $n$ nodes, the method generates initial node embeddings using the Node2Vec algorithm, which applies a random-walk-based graph embedding to the clique-expanded representation of the hypergraph[9]. Let the feature vector of a node $v_i$ be represented as $\mathbf{x}_i$. For a given hyperlink $e_p$, its feature vector $\mathbf{y}_p$ is computed by aggregating the embeddings of all associated nodes through mean pooling:

$$\mathbf{y}_p = \frac{1}{|e_p|} \sum_{v_i \in e_p} \mathbf{x}_i.$$

The model predicts the likelihood of $e_p$ being a valid hyperlink using a simple neural network layer with a sigmoid activation function, defined as:

$$S_p = \sigma\left(\mathbf{W}_{\text{score}}\mathbf{y}_p + \mathbf{b}_{\text{score}}\right),$$

where $\mathbf{W}_{\text{score}}$ and $\mathbf{b}_{\text{score}}$ are trainable parameters. The resulting score $S_p \in [0, 1]$ serves as a probabilistic indicator of the hyperlink. While NVM provides a straightforward approach, transforming a hypergraph into its clique-expanded form can lead to the loss of higher-order relationships. Moreover, the computational demands of Node2Vec grow significantly with the size of the graph, making the method less practical for large-scale hypergraphs.

**The self-attention-based graph neural network for hypergraphs (Hyper-SAGNN)**[10] utilizes attention mechanisms to enhance node features for hyperlink prediction. Initially, node features are derived by processing the adjacency matrix $\mathbf{A}$ (adjusted to remove self-loops) through a one-layer neural network:

$$\mathbf{x}_i = \tanh\left(\mathbf{W}_{\text{enc}}\mathbf{a}_i + \mathbf{b}_{\text{enc}}\right),$$

where $\mathbf{a}_i$ represents the column of the adjacency matrix corresponding to node $v_i$, and $\mathbf{W}_{\text{enc}}$, $\mathbf{b}_{\text{enc}}$ are learnable parameters. For a given hyperlink $e_p$, Hyper-SAGNN refines the features of the nodes within $e_p$ using both static and dynamic mechanisms. The dynamic attention coefficient $\alpha_{ij}$ quantifies the influence of node $j$ on node $i$ within the same hyperlink $e_p$, where both $i$ and $j$ are indices of nodes in $e_p$, and $j \neq i$. These coefficients are computed as:

$$\alpha_{ij} = \frac{\exp\left(\left(\mathbf{W}_i\mathbf{x}_i\right)^\top\left(\mathbf{W}_j\mathbf{x}_j\right)\right)}{\sum_{v_k\in e_p, k\neq i}\exp\left(\left(\mathbf{W}_i\mathbf{x}_i\right)^\top\left(\mathbf{W}_k\mathbf{x}_k\right)\right)},$$

where $\mathbf{W}_i$ and $\mathbf{W}_j$ are learnable parameters.

The refined feature vector for the hyperlink $e_p$ is obtained through a pooling operation that combines static and dynamic node features:

$$\mathbf{y}_p = \frac{1}{|e_p|}\sum_{v_i\in e_p}\left(\mathbf{s}_i - \mathbf{d}_i\right)^2,$$

where $\mathbf{s}_i$ and $\mathbf{d}_i$ represent the static and dynamic feature updates for node $i$, respectively, and the Hadamard power emphasizes their differences. Hyper-SAGNN is well-suited for capturing high-order structural information within hypergraphs but struggles with performance on sparse hypergraphs, such as metabolic networks.

**The Neural Hyperlink Predictor (NHP)**[11] shares a foundational similarity with Hyper-SAGNN but introduces a novel pooling mechanism that adapts to specific tasks by dynamically learning weights and integrating prior knowledge about the nodes. As with Node2Vec-mean (NVM), NHP begins by utilizing Node2Vec on the clique-expanded graph to initialize the feature vectors of the nodes. Let the feature vector of a node $v_i$ be denoted as $\mathbf{x}_i$. To refine these features, NHP applies a traditional graph neural network over each clique corresponding to hyperlinks in the original hypergraph. The updated feature vector for $v_i$ is calculated as follows:

$$\tilde{\mathbf{x}}_i = \mathrm{ReLU}\left(\mathbf{W}_{\text{conv1}}\mathbf{x}_i + \sum_{v_j, v_j \neq v_i \in e_p} \mathbf{W}_{\text{conv2}}\mathbf{x}_j\right),$$

where $\mathbf{W}_{\text{conv1}}$ and $\mathbf{W}_{\text{conv2}}$ are learnable weight matrices within the graph neural network, and ReLU serves as the default activation function. Once the node features are refined, NHP computes the feature vector for a hyperlink $e_p$ using a maximum-minimum pooling operation, designed to capture task-specific information by aggregating node-level features. This operation is defined as:

$$\mathbf{y}_p^{(\text{maxmin})_j} = \max_{v_i \in e_p} (\tilde{\mathbf{x}}_i)_j - \min_{v_i \in e_p} (\tilde{\mathbf{x}}_i)_j, \quad \text{for } j = 1, 2, \ldots, d_{\text{conv}},$$

where $d_{\text{conv}}$ represents the dimensionality of the node feature space in the graph neural network.

While NHP retains similarities with Node2Vec-mean in its reliance on Node2Vec for feature initialization, it mitigates limitations by leveraging its advanced pooling strategy. However, using the clique-expanded graph still risks losing high-order structural information, which can affect performance on larger or more complex datasets.

**Chebyshev Spectral Hyperlink predictor (CHESHIRE)**[12] is a deep learning framework designed to predict missing reactions in GEMs by leveraging the topological features of their metabolic networks. It models each metabolic network as a hypergraph, where metabolites are nodes and reactions are hyperlinks. Positive reactions correspond to existing ones, while negative reactions are synthetic and used for balancing during training. The method consists of four steps: feature initialization, feature refinement, pooling, and scoring. First, node features are initialized from the hypergraph's incidence matrix $\mathbf{H}$ using a one-layer neural network:

$$\mathbf{x}_i = \text{hard-tanh}\left(\mathbf{W}_{\text{enc}}\mathbf{h}_i + \mathbf{b}_{\text{enc}}\right),$$

where $\mathbf{W}_{\text{enc}}$ and $\mathbf{b}_{\text{enc}}$ are trainable parameters, and hard-tanh is a bounded activation function. In the refinement step, CHESHIRE applies a Chebyshev spectral graph convolutional network (CSGCN)[13] to enhance node features by incorporating local and global information. The refined features $\tilde{\mathbf{x}}_i$ are computed recursively as:

$$\tilde{\mathbf{x}}_i = \text{hard-tanh}\left(\sum_{k=1}^{K} \mathbf{W}_{\text{conv}}^{(k)}\mathbf{Z}_i^{(k)}\right),$$

where $\mathbf{Z}_i^{(k)}$ are recursively defined using Chebyshev polynomials:

$$\mathbf{Z}_i^{(1)} = \mathbf{x}_i, \quad \mathbf{Z}_i^{(2)} = \mathbf{L}\mathbf{Z}_i^{(1)}, \quad \mathbf{Z}_i^{(k)} = 2\mathbf{L}\mathbf{Z}_i^{(k-1)} - \mathbf{Z}_i^{(k-2)}.$$

Here, $\mathbf{L}$ is the scaled normalized Laplacian matrix of the graph.

To aggregate node features into reaction-level features, CHESHIRE combines Frobenius norm-based pooling and maximum-minimum pooling. Finally, the pooled features are passed into a neural network to predict the probability of a reaction's existence:

$$S_p = \sigma\left(\mathbf{W}_{\text{score}}\mathbf{y}_p + \mathbf{b}_{\text{score}}\right),$$

where $S_p$ represents the confidence score for each reaction. By integrating a CSGCN for feature refinement and combining complementary pooling methods, CHESHIRE effectively captures complex structural relationships in metabolic networks.

**Hypergraph Convolution network and attention mechanism integrated explorer for gaps prediction of metabolism (CLOSEgaps)**[14] is a deep learning framework designed to predict missing reactions in GEMs. The method consists of five main steps: mapping GEMs to a hypergraph, negative reaction sampling, feature initialization, feature refinement, and ranking or prediction. Initially, GEMs are represented as hypergraphs $\mathcal{H}$, where metabolites are hypernodes and reactions are hyperedges. Negative reactions are generated using metabolic network data and the ChEBI database, introducing synthetic reactions to balance the dataset. To initialize features, hypernode features $\mathbf{X}^{(0)}$ are computed by combining structural and biochemical information. This is achieved through a fully connected layer:

$$\mathbf{X}^{(0)} = \text{Cat}(\text{Linear}(\mathbf{H}_p), \text{Linear}(\mathbf{S})),$$

where $\mathbf{H}_p$ is the hypergraph with negative reactions, and $\mathbf{S}$ is a metabolite similarity matrix.

Features are refined using a multi-channel hypergraph convolutional network (HGCN) with attention mechanisms, which incorporate both local and global structural information. For a given hypergraph $\mathcal{H}$, the embedding of a hypernode $v_i$ at the $(l+1)$-th layer is updated as:

$$\mathbf{x}_i^{(l+1)} = \sigma\left(\sum_{j=1}^{n}\sum_{\epsilon=1}^{2m} H_{i\epsilon}H_{j\epsilon}W_\epsilon \mathbf{x}_j^{(l)}\mathbf{P}\right),$$

where $H_{i\epsilon}$ represents the attention score between hypernode $v_i$ and hyperedge $\epsilon$, $W_\epsilon$ is the hyperedge weight, and $\sigma(\cdot)$ is a nonlinear activation function. The attention score $H_{i\epsilon}$ is computed using a similarity function to capture interactions between nodes and hyperedges:

$$H_{i\epsilon} = \frac{\exp\left(\sigma\left(\mathrm{sim}(\mathbf{x}_i^{(l)}\mathbf{P}, \mathbf{x}_\epsilon^{(l)}\mathbf{P})\right)\right)}{\sum_{k\in\mathcal{N}_i}\exp\left(\sigma\left(\mathrm{sim}(\mathbf{x}_i^{(l)}\mathbf{P}, \mathbf{x}_k^{(l)}\mathbf{P})\right)\right)}.$$

The refined features are then used to rank candidate reactions based on their confidence scores, enabling the prediction of missing reactions in GEMs. By integrating hypergraph convolution and attention mechanisms, CLOSEgaps effectively captures complex relationships within metabolic networks, providing a powerful tool for metabolic network reconstruction.

### Supplementary Note 2: Difference between MuSHIN and CLOSEgaps

Both CLOSEgaps[14] and MuSHIN aim to predict missing reactions in GEMs. However, while CLOSEgaps primarily focuses on topological and basic biochemical properties of metabolic networks, MuSHIN further integrates detailed chemical structure embeddings and advanced hypergraph-based learning to capture complex biochemical patterns and improve prediction performance.

1. In the feature initialization step, CLOSEgaps relies on structural information derived from the hypergraph's incidence matrix and basic metabolite similarity matrices. In contrast, MuSHIN

employs advanced chemical embeddings generated through transformer-based models, specifically RXNFP and ChemBERTa. These embeddings provide comprehensive chemical representations capturing molecular-level structural features of metabolites and reactions, enabling more nuanced biological understanding and richer input representations for subsequent learning stages.

2. During feature refinement, CLOSEgaps uses a multi-channel hypergraph convolutional network (HGCN) combined with conventional attention mechanisms. MuSHIN, however, adopts a sophisticated dynamic attention mechanism integrated within its hypergraph neural network (HGNN). This approach involves iterative feature updates via node-to-edge and edge-to-node attention, dynamically capturing the intricate, higher-order relationships and chemical interactions between metabolites and reactions throughout the network refinement process.

3. Additionally, MuSHIN integrates advanced deep learning methodologies such as graph normalization and alpha dropout techniques. Graph normalization stabilizes the learning process by reducing covariate shifts across hypergraph layers, while alpha dropout regularizes neural network training by randomly excluding neurons, thereby promoting robustness and generalization in the reaction prediction tasks.

By combining chemical embeddings and an attention-enhanced HGNN, MuSHIN significantly extends the capabilities of CLOSEgaps, enabling it to capture the structural and chemical complexities of metabolic networks more comprehensively.

### Supplementary Note 3: Internal validation

#### Prediction Workflow

MuSHIN combines structural features from metabolic networks with chemical characteristics to predict potential missing reactions in GEMs. For each specific GEM, comprehensive training is conducted to have the model completely learn the relationships and chemical information of every metabolite and reaction within the metabolic network, allowing it to better adapt to and analyze the network under study. The prediction process uses the refined node and hyperedge representations obtained during training to generate probability scores for each reaction through the learned relationships, indicating the likelihood that a given reaction will be biologically valid.

**Scoring Reactions** To predict the validity of each reaction, we use the refined embeddings of metabolites (nodes) and reactions (hyperedges) produced by our model. Given the incidence matrix $\mathbf{H}$, which encodes the connectivity between metabolites and reactions, the model first uses multiple layers of hypergraph convolution to iteratively refine the representations of each reaction. This process enables the model to integrate information from distant nodes in the hypergraph, capturing the complex interrelations among metabolites and reactions.

The hypergraph convolution updates metabolite and reaction features through iterative processing. For each layer $l$, the model refines the metabolite representations, then uses these updated features to further refine the reaction embeddings. Specifically, for each reaction $r$, the refined representation $\mathbf{h}_r^{(l)}$ is obtained through the hypergraph convolution as follows:

$$\mathbf{h}_r^{(l)} = \text{HypergraphConv}(\mathbf{Z}^{(l-1)}, \mathbf{H}, \mathbf{H}^\top), \quad l = 1, \ldots, L, \tag{1}$$

where $\mathbf{Z}^{(0)}$ denotes the initial input features of the metabolites, and $\mathbf{Z}^{(l)}$ represents the refined features after the $l$-th layer. The incidence matrix $\mathbf{H}$ captures the connectivity between metabolites and reactions. Through stacking multiple convolutional layers ($L > 1$), the model gradually integrates broader contextual information from the hypergraph, capturing high-order relationships that are difficult for simpler models to represent.

The final refined representation $\mathbf{h}_r^{(L)}$ is then passed through a fully connected layer and activated by a softmax function to produce a probability score $\hat{y}_r$, indicating the likelihood that the reaction is biologically valid:

$$\hat{y}_r = \text{Softmax}(\mathbf{W}_{\text{pred}} \cdot \mathbf{h}_r^{(L)} + \mathbf{b}), \tag{2}$$

where $\mathbf{W}_{\text{pred}}$ and $\mathbf{b}$ are learnable parameters of the prediction layer. The softmax function ensures that $\hat{y}_r$ falls within the range $[0, 1]$, where a higher score indicates a greater likelihood of the reaction being biologically plausible.

**Loss Function** The training process uses a cross-entropy loss function to effectively distinguish between biologically valid and invalid reactions. Given the set of positive samples (valid reactions) $\mathcal{E}_p$ and negative samples (biologically improbable reactions) $\mathcal{E}_n$, the cross-entropy loss is defined as:

$$\text{Loss} = -\frac{1}{|\mathcal{E}_p| + |\mathcal{E}_n|} \left( \sum_{e_i \in \mathcal{E}_p} \log(\hat{y}_i) + \sum_{e_i \in \mathcal{E}_n} \log(1 - \hat{y}_i) \right), \tag{3}$$

where $\hat{y}_i$ represents the predicted probability of reaction $e_i$ being valid. The positive sample weight $w_p$ encourages the model to assign higher scores to biologically valid reactions, while the negative sample weight $w_n$ ensures lower predicted scores for invalid reactions. By optimizing this loss function, the model adjusts its parameters to maximize its ability to distinguish between positive and negative samples.

During training, we use the Adam optimizer to update the model parameters, making the training process more stable and allowing faster convergence. We tuned hyperparameters such as learning rate, batch size, and the number of epochs to achieve optimal performance.

**Reaction Classification** In the prediction phase, the model evaluates the test set, which contains a mixture of known valid reactions and artificially generated negative reactions. For each reaction $r$, the model generates a probability score $\hat{y}_r$ that indicates the likelihood of the reaction being biologically valid. To determine the final classification of each reaction, we use a decision threshold $\tau = 0.5$. If $\hat{y}_r \geq \tau$, the reaction is classified as valid, otherwise, it is classified as invalid:

$$\text{Prediction}(r) = \begin{cases} \text{Valid}, & \text{if } \hat{y}_r \geq 0.5, \\ \text{Invalid}, & \text{if } \hat{y}_r < 0.5. \end{cases} \tag{4}$$

This threshold can be adjusted based on the performance on the validation set to better fit specific application requirements. The final predicted classes are then compared to the ground truth labels in the test set to calculate metrics such as the F1-score, providing a comprehensive evaluation of the model's ability to generalize to unseen biochemical reactions.

### Evaluation Metrics

To evaluate the predictive performance of MuSHIN, we use five core classification metrics: accuracy, F1 score, area under the receiver operating characteristic curve (AUROC), area under the precision-recall curve (AUPRC), recall, and precision. These metrics provide a comprehensive assessment of the model's ability to predict biologically valid reactions.

**Accuracy**: The proportion of correctly classified reactions, including both positive (valid) and negative (invalid), to the total number of reactions. It is defined as:

$$\text{Accuracy} = \frac{\text{TP} + \text{TN}}{\text{TP} + \text{TN} + \text{FP} + \text{FN}},$$

where TP, TN, FP, and FN represent true positives, true negatives, false positives, and false negatives, respectively.

**F1 Score**: The harmonic mean of precision and recall, capturing the balance between these two metrics. It is particularly useful for evaluating performance in datasets with class imbalance. The F1 score is computed as:

$$\text{F1} = 2 \cdot \frac{\text{Precision} \cdot \text{Recall}}{\text{Precision} + \text{Recall}}.$$

**AUROC (Area Under the ROC Curve)**: AUROC measures the model's ability to distinguish between valid and invalid reactions across various thresholds. A higher AUROC indicates better discrimination performance.

**AUPRC (Area Under the Precision-Recall Curve)**: AUPRC evaluates the model's performance in datasets with class imbalance. It focuses on the trade-off between precision and recall and is particularly useful when the proportion of valid reactions is significantly smaller than invalid reactions. A higher AUPRC indicates better performance in detecting valid reactions.

**Precision**: The proportion of correctly predicted valid reactions out of all reactions predicted as valid. Precision is defined as:

$$\text{Precision} = \frac{\text{TP}}{\text{TP} + \text{FP}}.$$

**Recall**: The proportion of correctly predicted valid reactions out of all actual valid reactions. Recall is defined as:

$$\text{Recall} = \frac{\text{TP}}{\text{TP} + \text{FN}}.$$

**Hyperparameter settings**

All models were trained for a maximum of 100 epochs, with the model from the epoch achieving the highest F1 score on the validation set selected as the final model. Both MuSHIN and CLOSEgaps used an embedding dimension of 64, a convolution dimension of 128, a learning rate of $10^{-2}$, a weight decay of $10^{-3}$, and a batch size of 256. The hypergraph neural network (HGNN) in MuSHIN consisted of 2 layers. For other baseline models, CHESHIRE and HGNN used a feature dimension of 256, with CHESHIRE's Chebyshev Spectral Graph Convolution Network (CSGCN) having a convolution dimension of 128. CHESHIRE was trained with a learning rate of $10^{-2}$. NHP used a feature dimension of 512 and a convolution dimension of 256. Both HGNN and NHP were trained with a learning rate of $10^{-3}$. The weight decay for all three baseline models (CHESHIRE, HGNN, and NHP) was set to $5 \times 10^{-4}$.

**Threshold scores**

In **Figure 2**, we applied a fixed threshold score of 0.5 to determine whether an unseen reaction should be classified as valid or invalid. However, model performance can vary depending on the thresholding strategy. To further examine this effect, we evaluated MuSHIN and all baseline models (HGNN, NHP, CHESHIRE, and CLOSEgaps) on 108 BiGG GEMs using two alternative thresholding approaches: setting the threshold as the mean or the median of all predicted scores for reactions in the test set. The results indicate that the overall performance distributions under the mean and median thresholds are highly similar to those obtained with the fixed 0.5 threshold, with the median threshold leading to slightly lower overall performance. Nonetheless, MuSHIN remains significantly superior to all baseline models across all evaluation metrics (**Supplementary Fig. 1**). Since AUROC and AUPR do not depend on the threshold choice, they were excluded from this comparison.

**Negative sampling strategies**

Negative sampling plays a crucial role in constructing training datasets for reaction prediction. Different strategies for selecting negative reactions can impact model performance. In our approach, we generate negative reactions by replacing a fraction ($\alpha$) of metabolites in known reactions with randomly selected metabolites from a global metabolite pool. This ensures that the generated negative reactions remain chemically invalid while preserving structural similarity to real metabolic reactions. In Figure 2, we used $\alpha = 0.5$ to create negative samples. To further analyze the sensitivity of different models to negative sampling strategies, we evaluated MuSHIN and all baseline methods (HGNN, NHP, CHESHIRE, and CLOSEgaps) on 108 BiGG GEMs using two additional settings: $\alpha = 0.3$ and $\alpha = 0.7$. Lower values of $\alpha$ make negative reactions easier to distinguish, whereas higher values generate reactions that are structurally closer to real ones, making classification more challenging. As expected, all baseline models performed worse across all evaluation metrics when $\alpha$ increased from 0.3 to 0.7 (**Supplementary Fig. 2 and 3**). In contrast, MuSHIN remained highly stable, showing minimal performance variation across different negative sampling difficulties. This demonstrates the robustness of MuSHIN in accurately distinguishing true metabolic reactions from chemically invalid ones, even when negative samples closely resemble real reactions.

**Negative sampling ratios**

The ratio of positive to negative reactions in negative sampling can also influence model performance. In **Figure 2**, we applied a 1:1 ratio, meaning each positive reaction was paired with one negative reaction. To further investigate this effect, we evaluated MuSHIN and all baseline methods (HGNN, NHP, CHESHIRE, and CLOSEgaps) on 108 BiGG GEMs under two alternative negative sampling ratios: 1:2 and 1:3, where each positive reaction was paired with two or three negative reactions, respectively. The results show that MuSHIN continues to significantly outperform all baseline models across all evaluation metrics (**Supplementary Fig. 4 and 5**). As the number of negative samples increases, baseline models experience a notable drop in F1 Score, Precision, and AUPRC, while MuSHIN maintains stable performance across all metrics. This robustness highlights MuSHIN's ability to generalize effectively even under challenging data conditions.

### Supplementary Note 4: External validation

**Generation of GEMs** All draft GEMs (**Supplementary Table 1**) were reconstructed using the standard CarveMe[15] pipeline. Only growth phenotypes were used for the built-in gap-filling algorithm. To fill the gaps in a given draft GEM, we first selected candidate BiGG reactions whose model likelihood scores ($\mathcal{S}_\mu$) were equal to or greater than 0.999. These candidates were then ranked according to their composite scores $\mathcal{S}(r)$ as defined in Equation 5, from highest to lowest.

The top 100 reactions were iteratively added to the draft GEM. For each added reaction, we tested whether it led to increased biomass flux, which would indicate the establishment of energy-generating cycles (EGCs). EGCs are thermodynamically infeasible cycles capable of charging energy molecules without nutrient consumption. We employed the method developed by Fritzemeier et al.[16] to detect EGCs. Specifically, we created 17 energy dissipation reactions and maximized the flux of one reaction at a time while prohibiting all influx into the model. These dissipation reactions correspond to 17 different types of energy metabolites: ATP, CTP, GTP, UTP, ITP, NADH, NADPH, $FADH_2$, $FMNH_2$, Q8H2, Mql8, Mql6, Mql7, 2Dmmql8, AcCoA, L-Glutamate, and proton.

Any non-zero flux through one of the 17 dissipation reactions indicates the presence of an EGC that can generate the associated energy metabolite. If the added reaction was reversible, we resolved detected EGCs by constraining its flux bounds: the flux was restricted to be non-positive or non-negative if the reaction had a positive or negative flux in the EGC test, respectively. We skipped the irreversible added reactions. For the fermentation metabolite test where bacteria were grown in anaerobic conditions, we skipped any reaction containing oxygen as a reactant or product. We also excluded reactions that increased biomass flux above the known maximum bacterial growth rate (2.81 $h^{-1}$, equivalent to 15 min/generation). This process continued until 100 reactions were successfully added.

**Culture media compositions** The culture media compositions used for growth simulations were determined to reproduce the experimental conditions under which phenotypes were measured. While the fermentation product test dataset originated from multiple experiments with potentially varying culture media, we followed the strategy described in Zimmermann et al.[17] and assumed that all experiments were performed under the same growth medium. We adopted the fermentation test medium composition and their maximally allowed fluxes developed in the same study (accessible from https://github.com/Waschina/gapseqEval).

For the amino acid secretion test, M9 minimal medium (with glucose) was used. Glucose had a maximum uptake rate of 10 mmol/gDW/h and all other compounds in the medium were unconstrained. For the substrate utilization test, GEMs were also constrained to the same M9 minimal medium, where the default sources of carbon, nitrogen, sulfur, and phosphorus were glucose, ammonia, sulfate, and phosphate, respectively. For *Shewanella oneidensis*, the default carbon source was DL-lactate. To simulate growth on each substrate in Biolog arrays, the default source with the same type as the substrate (i.e., carbon, nitrogen, sulfur, or phosphorus) in the M9 minimal medium was replaced with the test substrate. The maximum uptake rate for all Biolog substrates was 10 mmol/gDW/h and all other compounds in the

M9 medium were unconstrained. The M9 recipe was obtained from the CarveMe repository (https://github.com/cdanielmachado/carveme).

**Simulations of metabolic phenotypes** We adopted a similar strategy as used in Zimmermann et al.[17] to compute outflux values of secreted metabolites. For each GEM, we ran parsimonious Flux Balance Analysis (pFBA)[18] to avoid nutrient influxes that do not contribute to biomass and used the pFBA solution to constrain import fluxes. Flux variability analysis (FVA)[19] was applied to predict the maximum secretion fluxes of metabolites under the constraint of maximum growth rate. Metabolites with a normalized outflow (secretion flux divided by biomass) larger than $10^{-5}$ were considered as produced by the GEM. Therefore, our algorithm classified each metabolite as being produced or not produced by the GEM, which could be directly compared to the observed data. We examined the ability of draft GEMs and their gap-filled versions to produce a comprehensive list of 236 metabolites with BiGG IDs, as originally published by Zimmermann et al.[17]. We used Flux Balance Analysis (FBA)[20] to simulate bacterial growth on each substrate in Biolog phenotype arrays. The medium for each substrate was developed using the approach described in Culture media compositions. A growth phenotype was considered positive if the growth rate was at least 0.01 $h^{-1}$.

**Causal reaction inference** For each metabolite secreted by the gap-filled but not the draft GEM, we employed Mixed Integer Linear Programming (MILP) to determine the minimal set of reactions that must be added during the gap-filling process to enable the experimentally observed phenotype.

We modeled the flux activity of each candidate reaction using a binary variable $A$, subject to two linear constraints: (1) $f - f_{\text{min}} \cdot A \geq 0$ and (2) $f - f_{\text{max}} \cdot A \leq 0$, where $f$ denotes the reaction flux, and $f_{\text{min}}$ and $f_{\text{max}}$ represent the lower and upper flux bounds, respectively. For exchange reactions, these bounds were set to $-1000$ and $1000$, establishing the default flux range. Under this formulation, when $A = 1$, the reaction operates with unconstrained flux within the interval $f \in [-1000, 1000]$, while $A = 0$ enforces zero flux ($f = 0$).

The optimization objective minimized the sum of all binary indicator variables, constrained by the requirement that the target metabolite secretion flux exceeds a positive threshold of $0.1$. The resulting minimal sum represents the minimum number of reactions required to gap-fill the draft GEM for successful metabolite production, thereby enabling identification of the essential reactions responsible for the observed phenotype.

**Network-based Feature Calculation and Selection** For each candidate reaction, we calculated four key features to evaluate its potential contribution to the draft GEM:

For each candidate reaction, we calculated four key features to evaluate its potential contribution to the draft GEM: (1) Connectivity, the fraction of metabolites in the candidate reaction that are already present in the draft model, which captures how well the reaction integrates with the existing metabolic network; (2) Model Likelihood, the prediction score from the MuSHIN HyperGraph Neural Network, representing the probability that the reaction is a correct addition to the GEM, with scores ranging from 0 to 1; (3) Similarity Score, the maximum cosine similarity between the candidate reaction and reactions already present in the draft model, calculated using stoichiometric vectors, where lower similarity indicates more novel reactions that could expand the metabolic capabilities of the model; and (4) Hub Metabolite Count, the number of metabolites in the reaction that participate in more than 5 reactions in the universal reaction network, as hub metabolites often represent central compounds in metabolism and reactions containing them are more likely to connect disparate parts of the network.

To ensure high-quality gap-filling, we first filtered candidate reactions to include only those with model likelihood scores of at least 0.999. Each of the four features was then normalized to a 0-1 scale using min-max normalization. For the similarity score, we applied an inverse transformation so that reactions with lower similarity (more novel reactions) received higher normalized scores. When the connectivity calculation involved an empty denominator, we assigned a default value of 0.5.

The normalized features were combined into a composite score using the following balanced weighting scheme:

$$\mathcal{S}(r) = 0.3\,\tilde{C} + 0.3\,\tilde{S}_{\text{pred}} + 0.2\,(1 - \tilde{S}_{\text{sim}}) + 0.2\,\tilde{N}_{\text{hub}} \tag{5}$$

where tildes denote normalized values. This weighting scheme equally prioritizes connectivity and model likelihood (30% each), while considering reaction novelty through similarity scores and metabolic network importance through hub metabolite counts (20% each). The top 100 reactions ranked by composite score were selected for gap-filling.

### Supplementary Note 5: Data and Resources

#### BiGG Models

For internal validation, we used a total of 108 genome-scale metabolic models (GEMs) obtained from the BiGG database (`http://BiGG.ucsd.edu`) as of January 2022. Before conducting gap-filling, we removed biomass reactions, exchange reactions, demand reactions, and sink reactions from each model, as these reaction types do not indicate missing knowledge within the metabolic network.

#### AGORA models

To ensure that our method generalizes across diverse GEM collections, we additionally included models from the AGORA database for internal validation. We used 818 genome-scale metabolic models (GEMs) from the AGORA collection. These models were downloaded from the Virtual Metabolic Human (VMH) database (`https://www.vmh.life`). The same preprocessing pipeline applied to the BiGG models was also used for the AGORA models to ensure compatibility and standardization across datasets.

#### Construction of the BiGG Universal Reaction Pool

The universal reaction pool was derived from the BiGG reaction database (`http://BiGG.ucsd.edu`). To ensure relevance to metabolic gap-filling, we excluded biomass, exchange, demand, and sink reactions. Additionally, reactions associated with compartments other than the cytosol, periplasm, and extracellular space were removed. Further filtering was applied to eliminate reactions lacking assigned names and those exhibiting imbalanced carbon stoichiometry (e.g.,

FPGS_tm and SUCptsp_l). Reactions with identifiers beginning with "r" followed by numerical digits were also excluded, as these are associated with non-microbial metabolic models. After curation, the final dataset included 10,393 unique metabolites and 16,337 unique reactions.

**Retrieval of SMILES Representations**

To obtain SMILES representations for metabolites, we used different strategies for BiGG and AGORA GEMs. For BiGG models, we retrieved SMILES from the BiGG database's metabolite information (http://bigg.ucsd.edu/data_access), which provides direct access to curated SMILES annotations. For AGORA GEMs, the metabolite names differ from those in BiGG, requiring a more extensive search process. We obtained AGORA models from the Virtual Metabolic Human (VMH) database (https://www.vmh.life/#downloadview) under the microbe collections. Metabolite annotations were extracted from each model's XML file, which contains cross-references to multiple external databases. Using these annotations, we queried KEGG (http://rest.kegg.jp), HMDB (http://www.hmdb.ca/metabolites), PubChem (https://pubchem.ncbi.nlm.nih.gov), and ChEBI (https://www.ebi.ac.uk/chebi) via their respective APIs to retrieve corresponding SMILES structures. The integration of these four complementary databases allowed us to recover SMILES for the majority of metabolites across AGORA GEMs.

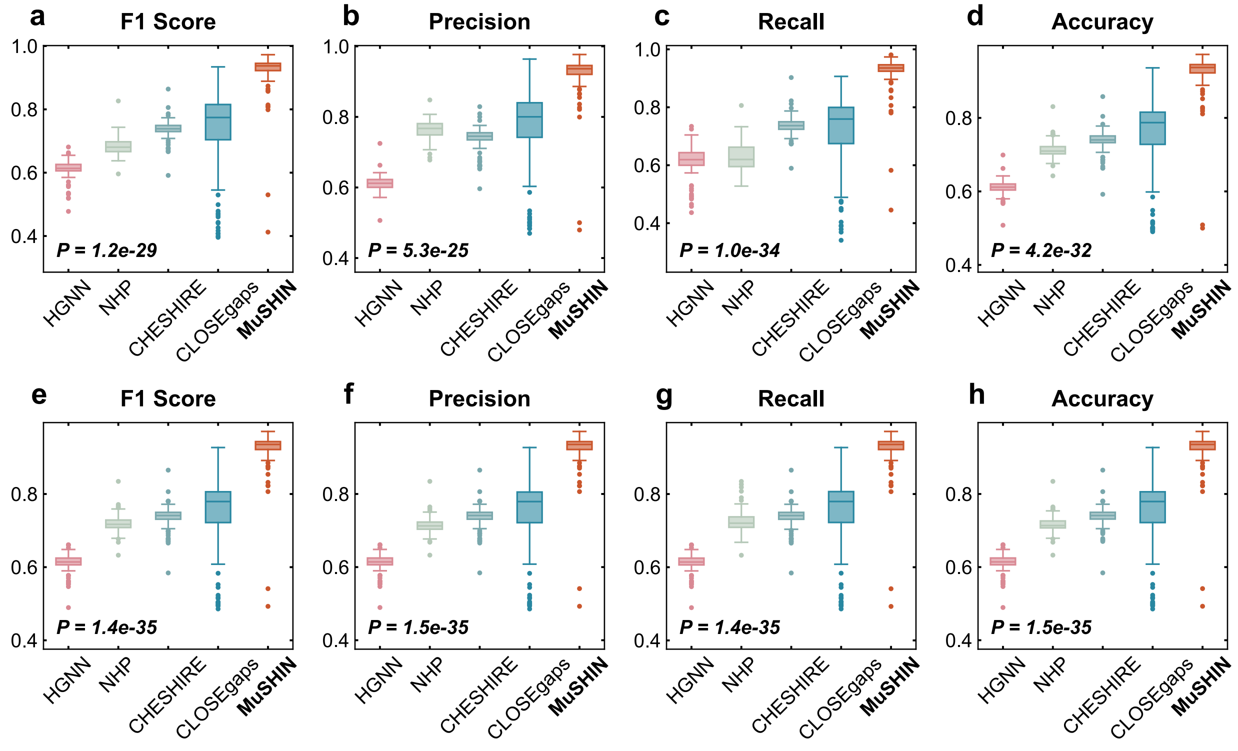


**Supplementary Fig. 1**: **Internal validation using artificially introduced gaps: Performance comparison of different thresholding methods. a-d** Boxplots of the performance metrics (F1 Score, Precision, Recall, and Accuracy) calculated on 108 BiGG GEMs using the mean threshold score, where the threshold is set as the mean predicted score of all reactions in the test set. **e-h** Boxplots of the same metrics calculated using the median threshold score, where the threshold is set as the median predicted score of all reactions in the test set. Each dot represents the performance on an individual GEM, with each data point representing the mean over 10 Monte Carlo runs to ensure robustness. Two-sided paired-sample t-tests were conducted between MuSHIN and the second-best baseline; exact p-values are reported. Source data are provided as a Source Data file.

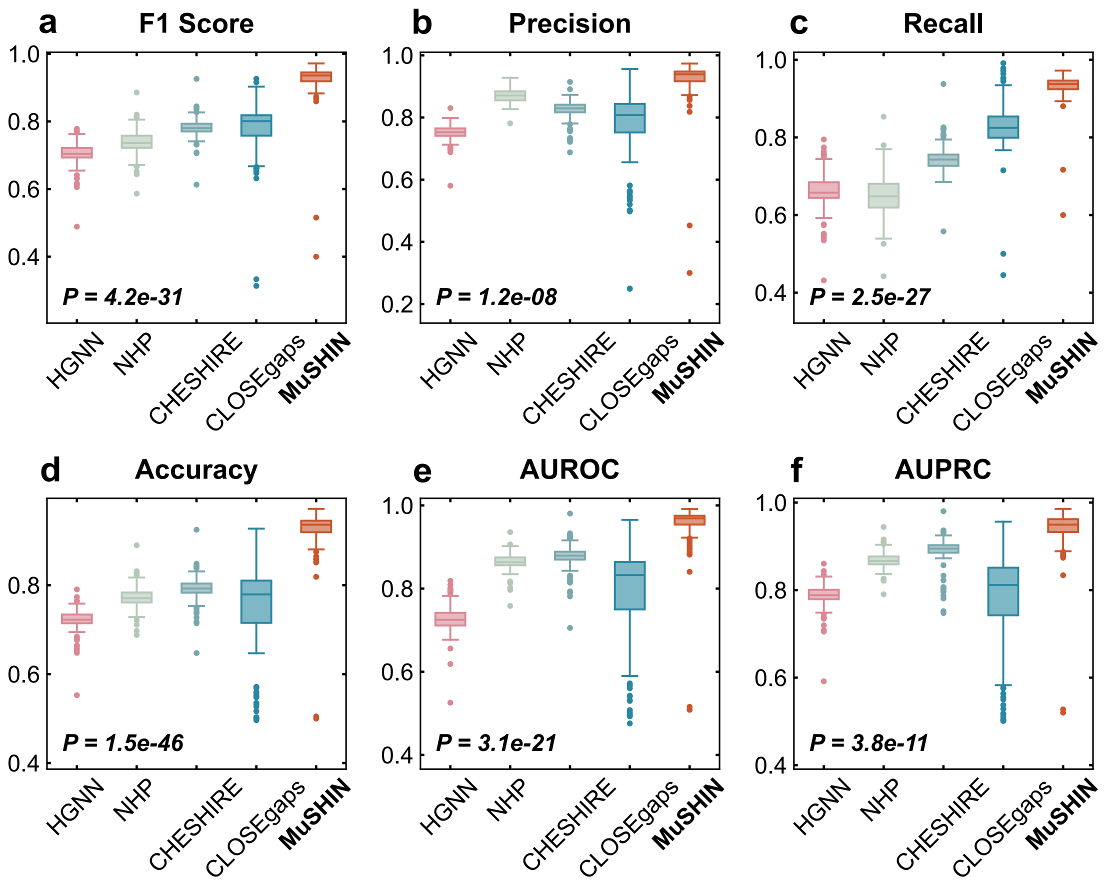


**Supplementary Fig. 2**: **Internal validation using artificially introduced gaps: Performance comparison under negative sampling with 30% metabolite replacement. a-f** Boxplots of the performance metrics (F1 Score, Precision, Recall, Accuracy, AUROC, and AUPRC) calculated on 108 BiGG GEMs using negative sampling where 30% of metabolites in each reaction were replaced. Each dot represents the performance on an individual GEM, with each data point representing the mean over 10 Monte Carlo runs to ensure robustness. Two-sided paired-sample t-tests were conducted between MuSHIN and the second-best baseline; exact p-values are reported. Source data are provided as a Source Data file.

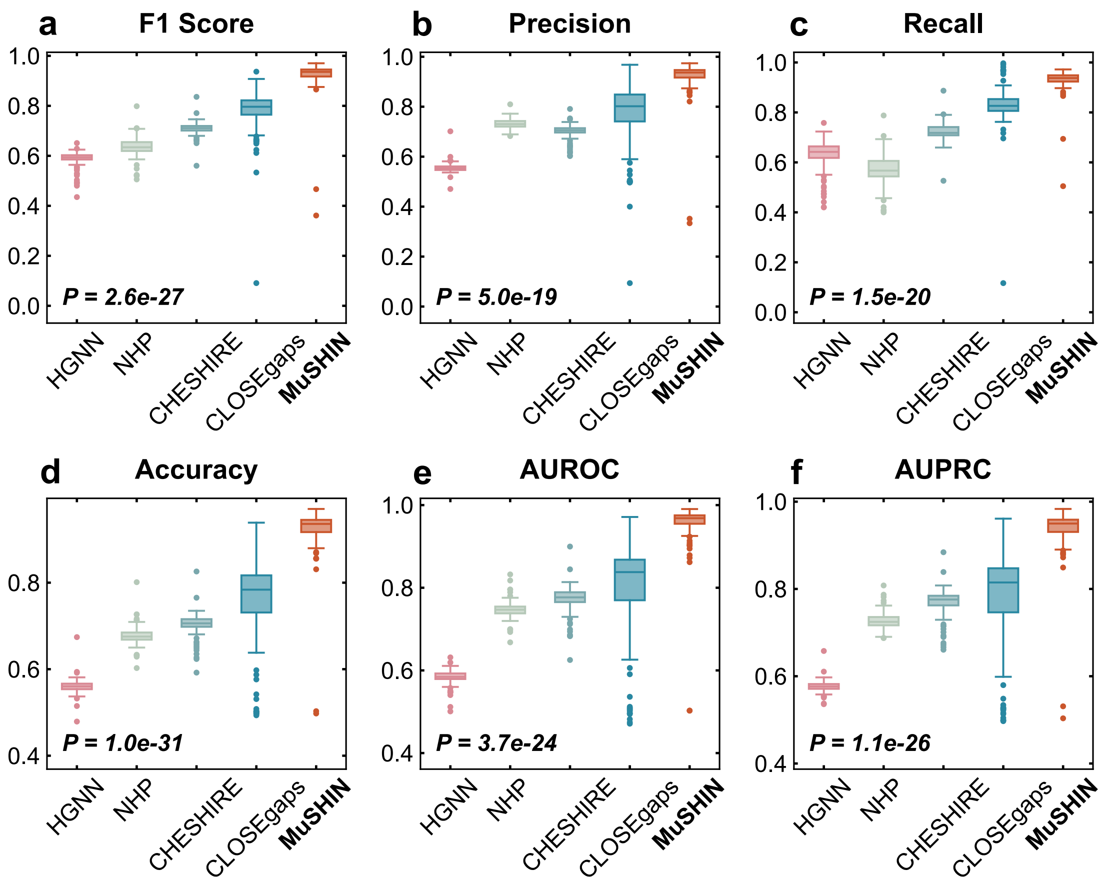


**Supplementary Fig. 3**: **Internal validation using artificially introduced gaps: Performance comparison under negative sampling with 70% metabolite replacement. a-f** Boxplots of the performance metrics (F1 Score, Precision, Recall, Accuracy, AUROC, and AUPRC) calculated on 108 BiGG GEMs using negative sampling where 70% of metabolites in each reaction were replaced. Each dot represents the performance on an individual GEM, with each data point representing the mean over 10 Monte Carlo runs to ensure robustness. Two-sided paired-sample t-tests were conducted between MuSHIN and the second-best baseline; exact p-values are reported. Source data are provided as a Source Data file.

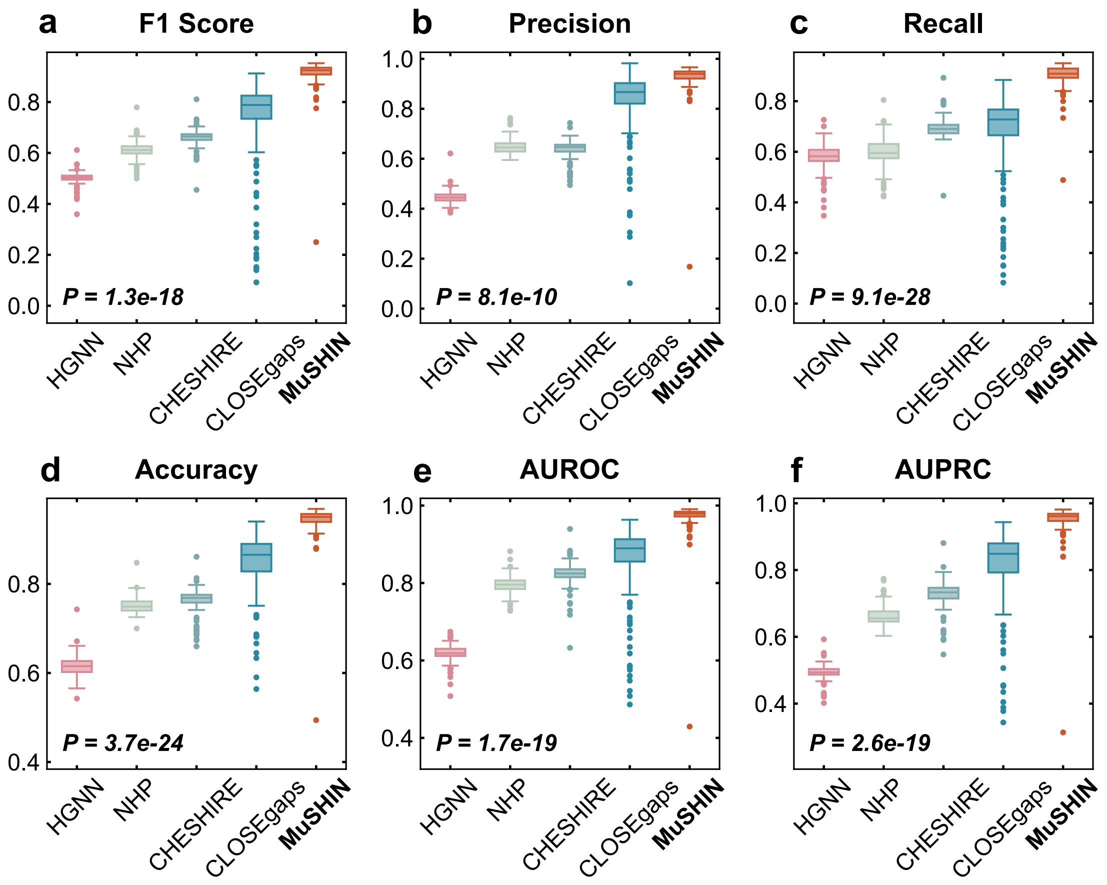


**Supplementary Fig. 4**: **Internal validation using artificially introduced gaps: Performance comparison under a negative sampling ratio of 1:2. a-f** Boxplots of the performance metrics (F1 Score, Precision, Recall, Accuracy, AUROC, and AUPRC) calculated on 108 BiGG GEMs using a negative sampling ratio of 1:2, where each positive reaction is paired with two negative reactions. Each dot represents the performance on an individual GEM, with each data point representing the mean over 10 Monte Carlo runs to ensure robustness. Two-sided paired-sample t-tests were conducted between MuSHIN and the second-best baseline; exact p-values are reported. Source data are provided as a Source Data file.

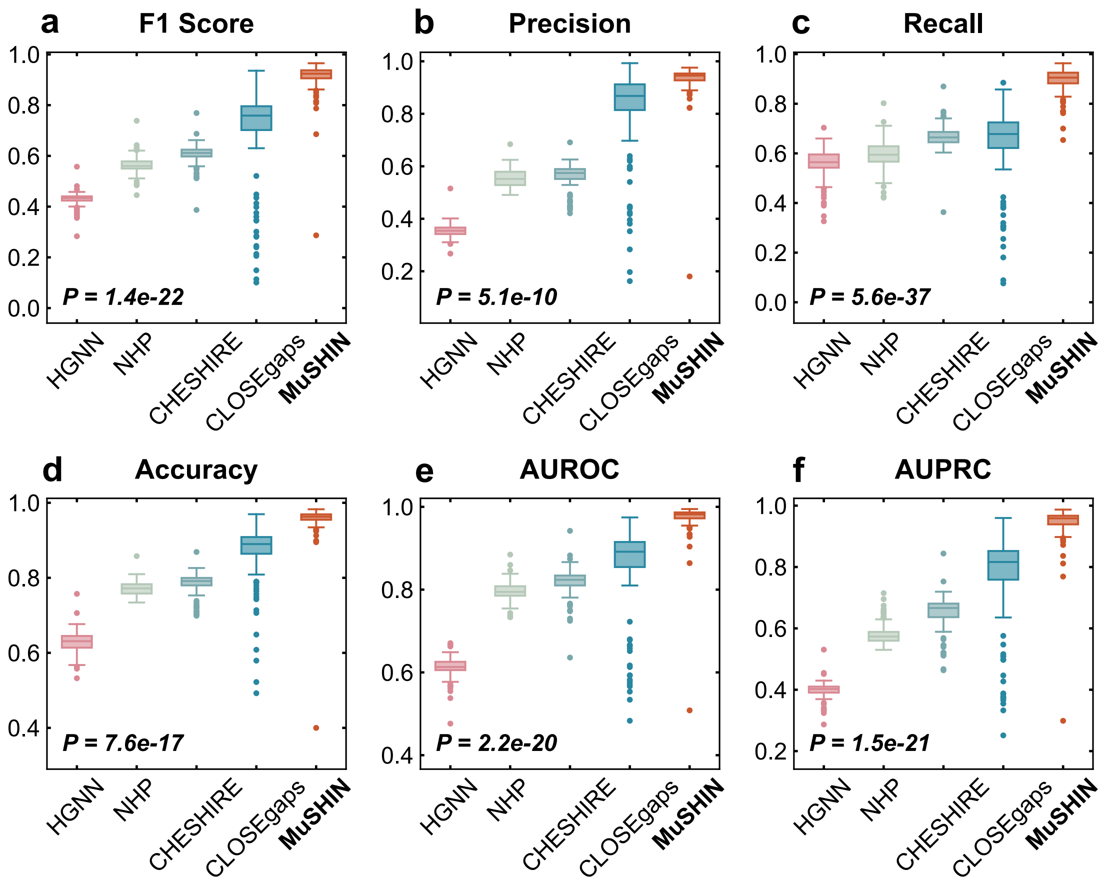


**Supplementary Fig. 5**: **Internal validation using artificially introduced gaps: Performance comparison under a negative sampling ratio of 1:3. a-f** Boxplots of the performance metrics (F1 Score, Precision, Recall, Accuracy, AUROC, and AUPRC) calculated on 108 BiGG GEMs using a negative sampling ratio of 1:3, where each positive reaction is paired with three negative reactions. Each dot represents the performance on an individual GEM, with each data point representing the mean over 10 Monte Carlo runs to ensure robustness. Two-sided paired-sample t-tests were conducted between MuSHIN and the second-best baseline; exact p-values are reported. Source data are provided as a Source Data file.

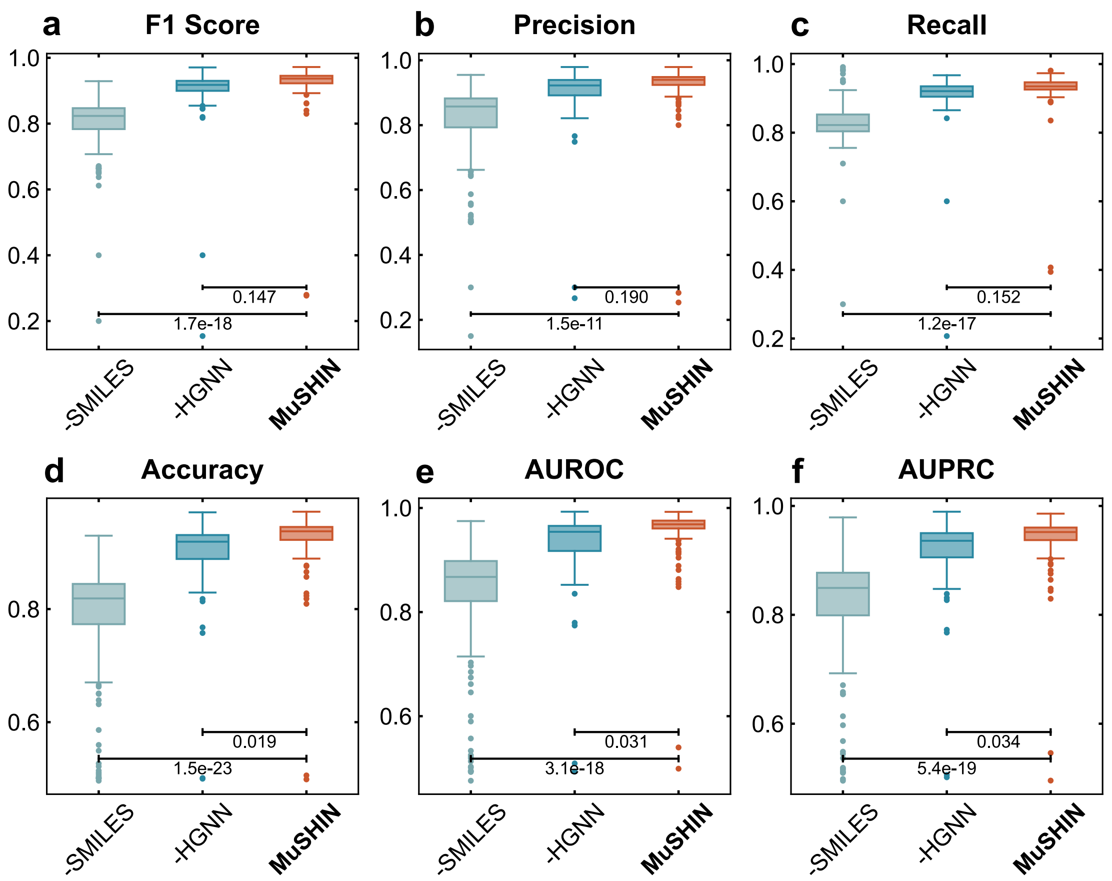


**Supplementary Fig. 6**: **Internal validation using artificially introduced gaps: Ablation study on MuSHIN components. a-f** Boxplots of the performance metrics (F1 Score, Precision, Recall, Accuracy, AUROC, and AUPRC) calculated on 108 BiGG GEMs for different ablated versions of MuSHIN. Variants include removing the multi-attention HGNN component (-HGNN), replacing SMILES embeddings with molecular fingerprints (-SMILES). Each dot represents the performance on an individual GEM, with each data point representing the mean over 10 Monte Carlo runs to ensure robustness. Source data are provided as a Source Data file.

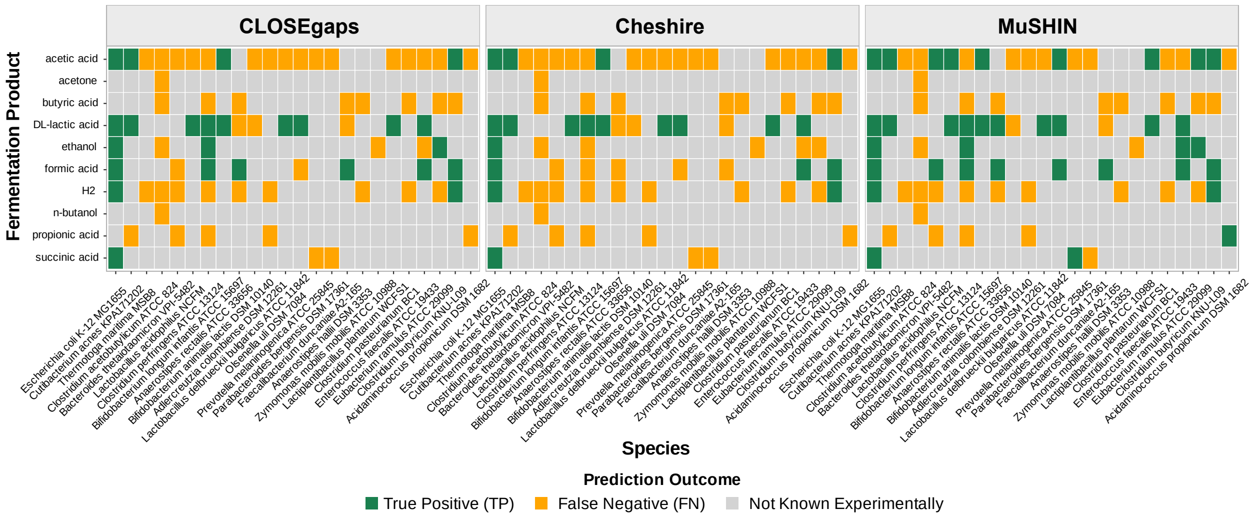


**Supplementary Fig. 7**: **Performance comparison of gap-filling algorithms for predicting fermentation products with 100 candidate reactions.** Heatmaps compare the performance of three algorithms (MuSHIN, Cheshire, and CLOSEgaps) in predicting fermentation products across 24 bacterial strains. Cell color indicates the outcome: green (True Positive), orange (False Negative), or grey (not known experimentally).

Supplementary Table 1: Bacterial genomes used in our external validation for testing fermentation products.

| **NCBI Assembly** | **Taxonomy** |
|---|---|
| GCF_000005845.2 | *Escherichia coli* str. K-12 substr. MG1655 |
| GCF_000008345.1 | *Cutibacterium acnes* KPA171202 |
| GCF_000008545.1 | *Thermotoga maritima* MSB8 |
| GCF_000008765.1 | *Clostridium acetobutylicum* ATCC 824 |
| GCF_000011065.1 | *Bacteroides thetaiotaomicron* VPI-5482 |
| GCF_000011985.1 | *Lactobacillus acidophilus* NCFM |
| GCF_000013285.1 | *Clostridium perfringens* ATCC 13124 |
| GCF_000020425.1 | *Bifidobacterium longum* subsp. *infantis* ATCC 15697 |
| GCF_000020605.1 | *Eubacterium rectale* ATCC 33656 |
| GCF_000022965.1 | *Bifidobacterium animalis* subsp. *lactis* DSM 10140 |
| GCF_000025885.1 | *Aminobacterium colombiense* DSM 12261 |
| GCF_000056065.1 | *Lactobacillus delbrueckii* subsp. *bulgaricus* ATCC 11842 |
| GCF_000143845.1 | *Olsenella uli* DSM 7084 |
| GCF_000144405.1 | *Prevotella melaninogenica* ATCC 25845 |
| GCF_000160535.1 | *Prevotella bergensis* DSM 17361 |
| GCF_000173975.1 | *Anaerobutyricum hallii* DSM 3353 |
| GCF_000175255.2 | *Zymomonas mobilis* subsp. *mobilis* ATCC 10988 |
| GCF_000389635.1 | *Clostridium pasteurianum* BC1 |
| GCF_000392875.1 | *Enterococcus faecalis* ATCC 19433 |
| GCF_000469345.1 | *Eubacterium ramulus* ATCC 29099 |
| GCF_001456065.2 | *Clostridium butyricum* KNU-L09 |
| GCF_001561955.1 | *Anaerotignum propionicum* DSM 1682 |
| GCF_000162015.1 | *Faecalibacterium prausnitzii* A2-165 |
| GCF_000203855.3 | *Lactobacillus plantarum* WCFS1 |

Supplementary Table 2: Unique correct fermentation product predictions by MuSHIN_100 algorithm compared to baseline (cheshire_100).

| NCBI Assembly | Unique Correct Products by MuSHIN_100 | Count |
|---|---|---|
| GCF_000011065.1 | acetic acid, formic acid | 2 |
| GCF_000013285.1 | ethanol, formic acid | 2 |
| GCF_000020605.1 | formic acid, DL-lactic acid | 2 |
| GCF_000143845.1 | acetic acid, formic acid | 2 |
| GCF_000469345.1 | acetic acid, ethanol | 2 |
| GCF_000011985.1 | acetic acid | 1 |
| GCF_000144405.1 | succinic acid | 1 |
| GCF_000162015.1 | formic acid | 1 |
| GCF_000203855.3 | acetic acid | 1 |
| GCF_000392875.1 | ethanol | 1 |
| GCF_001561955.1 | propionic acid | 1 |

### Supplementary References


1. Jeffrey D Orth, Ines Thiele, and Bernhard Ø Palsson. What is flux balance analysis? *Nat Biotechnol*, 28(3):245–248, March 2010.

2. Vinay Satish Kumar, Madhukar S Dasika, and Costas D Maranas. Optimization based automated curation of metabolic reconstructions. *BMC Bioinformatics*, 8(1):212, June 2007.

3. Ines Thiele, Nikos Vlassis, and Ronan M T Fleming. fastGapFill: efficient gap filling in metabolic networks. *Bioinformatics*, 30(17):2529–2531, May 2014.

4. Minoru Kanehisa and Susumu Goto. Kegg: kyoto encyclopedia of genes and genomes. *Nucleic acids research*, 28(1):27–30, 2000.

5. Muhan Zhang, Zhicheng Cui, Tolutola Oyetunde, Yinjie Tang, and Yixin Chen. Recovering metabolic networks using a novel hyperlink prediction method, 2016.

6. Zachary A King, Justin Lu, Andreas Dräger, Philip Miller, Stephen Federowicz, Joshua A Lerman, Ali Ebrahim, Bernhard O Palsson, and Nathan E Lewis. Bigg models: A platform for integrating, standardizing and sharing genome-scale models. *Nucleic acids research*, 44(D1):D515–D522, 2016.

7. Muhan Zhang, Zhicheng Cui, Shali Jiang, and Yixin Chen. Beyond link prediction: Predicting hyperlinks in adjacency space. In *Proceedings of the AAAI conference on artificial intelligence*, volume 32, 2018.

8. Govind Sharma, Prasanna Patil, and M Narasimha Murty. C3mm: Clique-closure based hyperlink prediction. In *IJCAI*, volume 20, pages 3364–3370, 2020.

9. Aditya Grover and Jure Leskovec. node2vec: Scalable feature learning for networks, 2016.

10. Ruochi Zhang, Yuesong Zou, and Jian Ma. Hyper-sagnn: a self-attention based graph neural network for hypergraphs, 2019.

11. Naganand Yadati, Vikram Nitin, Madhav Nimishakavi, Prateek Yadav, Anand Louis, and Partha Talukdar. Nhp: Neural hypergraph link prediction. In *Proceedings of the 29th ACM international conference on information & knowledge management*, pages 1705–1714, 2020.

12. Can Chen, Chen Liao, and Yang-Yu Liu. Teasing out missing reactions in genome-scale metabolic networks through hypergraph learning. *Nature Communications*, 14(1):2375, 2023.

13. Michaël Defferrard, Xavier Bresson, and Pierre Vandergheynst. Convolutional neural networks on graphs with fast localized spectral filtering. In D. Lee, M. Sugiyama, U. Luxburg, I. Guyon, and R. Garnett, editors, *Advances in Neural Information Processing Systems*, volume 29. Curran Associates, Inc., 2016.

14. Xiaoyi Liu, Hongpeng Yang, Chengwei Ai, Ruihan Dong, Yijie Ding, Qianqian Yuan, Jijun Tang, and Fei Guo. A generalizable framework for unlocking missing reactions in genome-scale metabolic networks using deep learning. *arXiv preprint arXiv:2409.13259*, 2024.

15. Daniel Machado, Sergei Andrejev, Marco Tramontano, and Kiran R Patil. Fast automated reconstruction of genome-scale metabolic models for microbial species and communities. *Nucleic Acids Research*, 46(15):7542–7553, 2018.

16. Christian J. Fritzemeier, Daniel Hartleb, Balázs Szappanos, Balázs Papp, and Martin J. Lercher. Erroneous energy-generating cycles in published genome scale metabolic networks: identification and removal. *PLoS Computational Biology*, 13(4):e1005494, 2017.

17. Julius Zimmermann, Christoph Kaleta, and Silvio Waschina. gapseq: informed prediction of bacterial metabolic pathways and reconstruction of accurate metabolic models. *Genome Biology*, 22(1):1–35, 2021.

18. Nathan E Lewis, Kory K Hixson, Tom M Conrad, Joshua A Lerman, Pep Charusanti, Ashoka D Polpitiya, Joshua N Adkins, Gregory Schramm, Samuel O Purvine, Daniel Lopez-Ferrer, et al. Omic data from evolved *E. coli* are consistent with computed optimal growth from genome-scale models. *Molecular Systems Biology*, 6(1):390, 2010.

19. Radhakrishnan Mahadevan and Christopher H Schilling. The effects of alternate optimal solutions in constraint-based genome-scale metabolic models. *Metabolic Engineering*, 5(4):264–276, 2003.

20. Jeremy S Edwards and Bernhard Ø Palsson. Metabolic flux balance analysis and the in silico analysis of escherichia coli k-12 gene deletions. *BMC Bioinformatics*, 1(1):1, 2000.
